# Conserved long noncoding RNA *TILAM* promotes liver fibrosis through interaction with PML in hepatic stellate cells

**DOI:** 10.1101/2023.07.29.551032

**Authors:** Cheng Sun, Chan Zhou, Kaveh Daneshvar, Arcadia J. Kratkiewicz, Amel Ben Saad, Anja Hess, Jennifer Y. Chen, Joshua V. Pondick, Samuel R. York, Wenyang Li, Sean Moran, Stefan Gentile, Raza Ur Rahman, Zixiu Li, Robert Sparks, Tim Habboub, Byeong-Moo Kim, Michael Y. Choi, Silvia Affo, Robert F. Schwabe, Yury V. Popov, Alan C. Mullen

**Affiliations:** Division of Gastroenterology, University of Massachusetts Chan Medical School, Worcester, MA, USA; Population and Quantitative Health Sciences, University of Massachusetts Chan Medical School, Worcester, MA USA; Division of Gastroenterology, Massachusetts General Hospital, Harvard Medical School, Boston, MA, USA; Liver Center, Department of Medicine, University of California, San Francisco, CA, USA; Broad Institute, Cambridge, MA, USA; Department of Liver, Digestive System, and Metabolism, Institut d’Investigacions Biomèdiques August Pi i Sunyer, Barcelona, Spain.; Department of Medicine, College of Physicians and Surgeons, Columbia University, New York, NY, USA.; Division of Gastroenterology, Hepatology and Nutrition, Beth Israel Deaconess Medical Center, Harvard Medical School, Boston, MA, USA

## Abstract

**Background & Aims:** Fibrosis is the common endpoint for all forms of chronic liver injury, and progression of fibrosis leads to the development of end-stage liver disease. Activation of hepatic stellate cells (HSCs) and their transdifferentiation to myofibroblasts results in the accumulation of extracellular matrix (ECM) proteins that form the fibrotic scar. Long noncoding (lnc) RNAs regulate the activity of HSCs and may provide targets for fibrotic therapies.

**Methods:** We identified lncRNA *TILAM* as expressed near *COL1A1* in human HSCs and performed loss-of-function studies in human HSCs and liver organoids. Transcriptomic analyses of HSCs isolated from mice defined the murine ortholog of *TILAM*. We then generated *Tilam*-deficient GFP reporter mice and quantified fibrotic responses to carbon tetrachloride (CCl_4_) and choline-deficient L-amino acid defined high fat diet (CDA-HFD). Co-precipitation studies, mass spectrometry, and gene expression analyses identified protein partners of *TILAM*.

**Results:** *TILAM* is conserved between human and mouse HSCs and regulates expression of ECM proteins, including collagen. *Tilam* is selectively induced in HSCs during the development of fibrosis *in vivo*. In both male and female mice, loss of *Tilam* results in reduced fibrosis in the setting of CCl_4_ and CDA-HFD injury models. *TILAM* interacts with promyelocytic leukemia protein (PML) to stabilize PML protein levels and promote the fibrotic activity of HSCs.

**Conclusion:** *TILAM* is activated in HSCs and interacts with PML to drive the development of liver fibrosis. Depletion of *TILAM* may serve as a therapeutic approach to combat the development of end stage liver disease.

## Introduction

Liver fibrosis develops as a result of chronic liver injury and can lead to cirrhosis and liver failure if the underlying cause of injury is not alleviated. Significant advances have been achieved in treating viral hepatitis^1, 2^, but other sources of chronic injury, including metabolic dysfunction-associated steatotic liver disease (MASLD)^3^ and primary sclerosing cholangitis (PSC) have limited treatment options beyond liver transplantation^4, 5^. While fibrosis is understood to be the common endpoint in nearly every form of chronic liver injury, there are currently no approved therapies that specifically target this pathologic process.

HSC myofibroblasts are the primary source of ECM that forms the progressive scar in liver fibrosis^6–8^. As a result of chronic liver injury, HSCs are activated and transdifferentiate from quiescent HSCs into activated HSCs/myofibroblasts, identified by increased secretion of ECM proteins, loss of lipid droplets, and increased contractility^9^.

Liver fibrosis can be reversible, and attenuation has been observed upon successful treatment of the underlying cause of injury such as in hepatitis B and C^10, 11^. With removal of the source of injury, the number of HSCs is reduced through apoptosis, and remaining HSCs revert to a more inactive state characterized by changes in gene expression and re-accumulation of lipid droplets^12, 13^.

Long noncoding RNAs (lncRNAs) are increasingly recognized as regulators and markers of disease^14–16^. lncRNAs are greater than 200 nucleotides (nt) in length and share features with messenger (m) RNAs, including 3’ polyadenylation in the majority of transcripts^17, 18^. Unlike mRNAs, lncRNAs do not serve as templates for translation and tend to be expressed at lower copy numbers than mRNAs^17, 18^. lncRNAs have been linked to diverse functions that include regulating chromatin, gene expression, and mRNA stability, and mediate these functions through interactions with DNA, RNA, and/or protein^19–21^.

Liver fibrosis is influenced by lncRNAs. *lincRNA-p21*, *MEG3*, and *GAS5* are predominantly anti-fibrotic. These lncRNAs are reduced with progression of fibrosis, and their induction in HSCs is associated with decreased expression of collagen type I (COL1A1) and smooth muscle actin (ɑ-SMA), as well as suppression of HSC proliferation^22–24^. In contrast, lncRNA *H19* is induced in cholestatic liver fibrosis^25^ and facilitates the activation of primary HSCs^26–28^. *H19* is imprinted and linked to multiple functions in development and disease^29^, and *lincRNA-p21*, *MEG3*, and *GAS5* are all expressed in many different cell types and tissues^30–32^, suggesting that modulating expression of these lncRNAs will affect more than just HSCs and liver fibrosis.

Here we describe lncRNA *TILAM* (<u>T</u>GF-β induced lncRNA <u>a</u>ctivating <u>m</u>yofibroblasts), which both uniquely identifies HSC myofibroblasts and regulates a program of fibrotic gene expression. We define the human and mouse orthologs and demonstrate that depletion of *TILAM* in human and mouse HSCs and in human liver organoids is sufficient to reduce collagen expression. We also find that loss of *TILAM* protects against the development of fibrosis *in vivo*. This activity is mediated, at least in part, through interaction with PML, which coordinates with *TILAM* to regulate ECM gene expression to promote liver fibrosis.

## Materials and methods

### Animal studies

All mouse experiments were approved by the IACUC of the Massachusetts General Hospital (2017000074) or Columbia University (AC-AAAF7452). For *in vivo* fibrosis studies, mice received 40% carbon tetrachloride (CCl_4_) diluted in olive oil or olive oil control by oral gavage (100 ul total volume) three times a week for four weeks^33^. For the choline-deficient, L-amino acid-defined, high-fat diet (CDA-HFD) model, mice were fed CDA-HFD chow consisting of 60% kcal fat and 0.1% methionine or control chow for twelve weeks^34^. HSCs were sorted from Lrat-Cre mice crossed with ZsGreen Cre reporter mice as described^8, 35^ either before or after receiving CCl_4_ by intraperitoneal injections (0.5 ml/g, dissolved in corn oil at a ratio of 1:3) three times a week for four weeks.

### Generation of Tilam mouse

*Tilam^gfp/gfp^*mice were generated with genome editing on a C57BL/6 background and were back-crossed to C57BL/6 mice for six generations before performing *in vivo* experiments.

### Cell culture

Primary human HSCs and LX-2 cells were cultured as previously described for HSCs^36^. Primary mouse HSCs were isolated and cultured as previously described^37^.

### Human liver organoids

H1 human ESCs (WA01, NIH registration number 0043) were obtained from WiCell, and ESCRO approval for their use was received from UMass Chan Medical School and Massachusetts General Hospital. hESCs were cultured and differentiated into liver organoids as previously described^21, 38^.

### RNA-Seq

Primary human HSCs were transfected in triplicate with siRNAs or locked nucleic acids (LNAs) and respective controls. Human HSC libraries were created using the New England Biolabs Directional Ultra II RNA Library Prep Kit prior to 150 nt paired-end sequencing on a HiSeq4000. Murine HSC libraries were created from sorted HSCs and prepared with the Illumina Truseq mRNA Stranded Prep Kit prior to 100 nt paired-end sequencing on a HiSeq2500.

### Mass spectrometry and immunoprecipitation

LX-2 cells were infected with lentivirus expressing *TILAM* and a scrambled version of *TILAM* (*sc-TILAM*), each fused to an aptamer that binds streptavidin (4x-S1m)^21^. Following streptavidin precipitation and RNase digestion, gel electrophoresis was performed, and proteins from 30-350 kD were analyzed by mass spectrometry.

A detailed description of methods is provided in the Supplemental Material.

## Results

### Identification of a human lncRNA near COL1A1

We evaluated RNA-seq data from human HSCs^39^ and identified an lncRNA located within a super-enhancer shared with *COL1A1* and contained in a co-expression network enriched for ECM genes^39^. The transcription start site (TSS) of the lncRNA is approximately 7.3 kb upstream of the TSS of *COL1A1,* and the gene is transcribed in the antisense direction compared to *COL1A1* (Figure 1*A*). There are separate regions of accessibility around the TSSs for *COL1A1* and the lncRNA, suggesting that each gene is under control of individual promoters (Supplementary Figure 1*A*).

**Figure 1.**
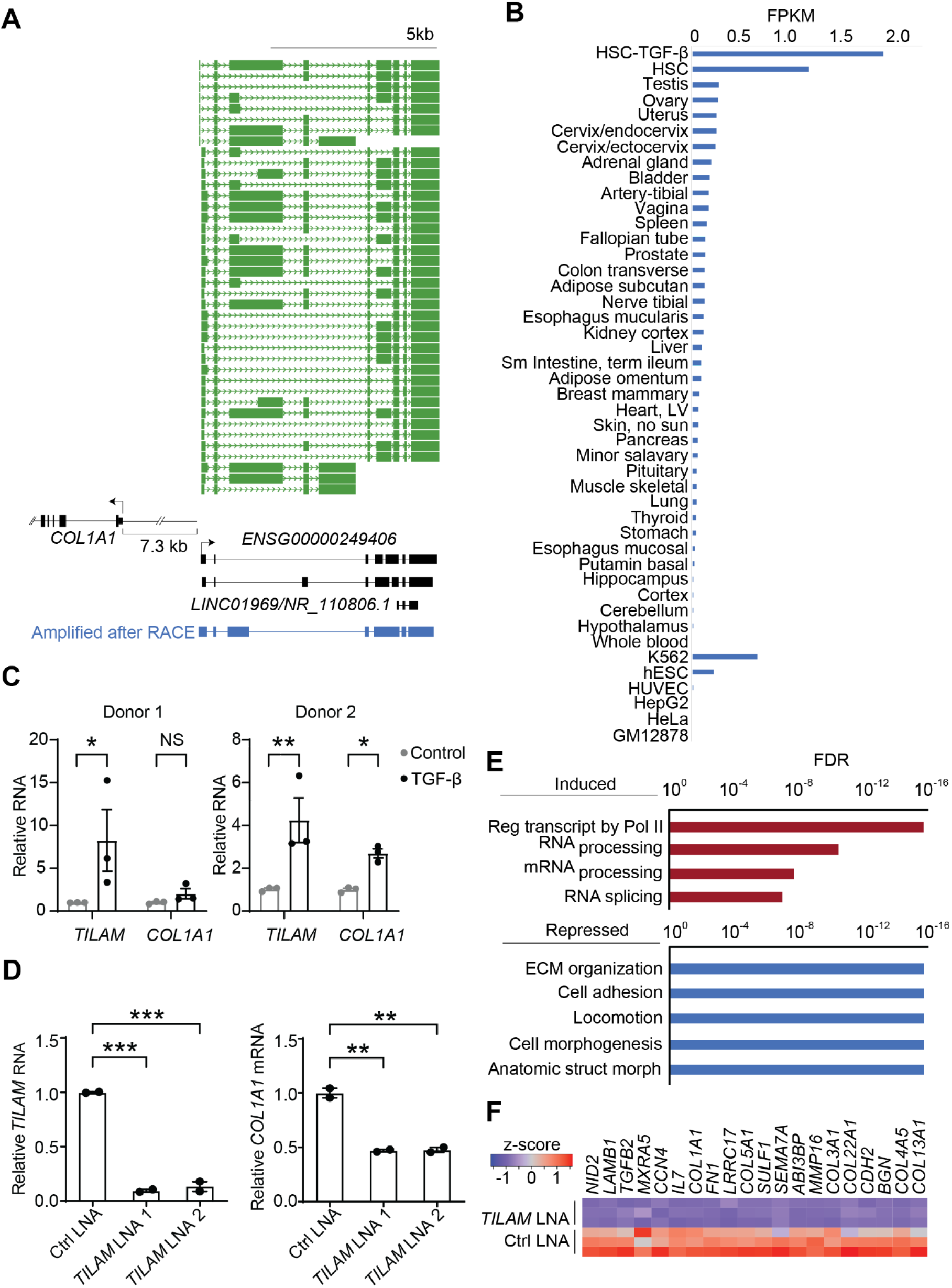
*TILAM* regulates the fibrotic activity of human HSCs. (*A*) The intron-exon structure of *ENSG00000249406* and *LINC01969* (GRCh38) are shown compared to the product cloned from primary human HSCs after RACE (Amplified after RACE). The distance between the TSS of *COL1A1* and *TILAM* is indicated. The isoforms previously annotated in HSCs^39^ are indicated in green. (*B*) *TILAM* gene expression was quantified across tissues and cell lines using previously published datasets^39^. (*C*) *TILAM* and *COL1A1* RNA levels were quantified by qRT-PCR in two primary human HSC lines in response to TGF-β treatment (5 ng/ml). * indicates p<0.05, ** indicates p<0.01, ns indicates p>0.05 (2-tailed unpaired *t* test). Error bars represent mean ± SEM. (*D*) *TILAM* was depleted in two primary human HSC lines using two different locked nucleic acid antisense oligonucleotides (LNA 1, LNA 2) compared to control LNA (Ctrl). *TILAM* (left) and *COL1A1* expression (right) were quantified by qRT-PCR. ** indicates p<0.01, *** indicates p<0.001 (one-way ANOVA with Dunnett’s multiple comparison). (*E*) *TILAM* was depleted in primary human HSCs (LNA 1), and RNA-seq was performed to compare gene expression to control LNA conditions (in triplicate). The GO categories representing the most induced (top, red) and repressed (bottom, blue) genes with depletion of *TILAM* are shown. False discovery rate (FDR) is indicated on the x-axis. (*F*) Heatmap displays expression of the 20 most repressed genes in the GO category of ECM organization with *TILAM* depletion.

Two isoforms of *TILAM* are annotated in GENCODE v43, and a shorter transcript containing the terminal three exons is also annotated separately (Figure 1*A*, bottom). RNA-seq analysis of human HSCs also identified additional isoforms in HSCs^39^ (Figure 1*A*, green tracks). We confirmed that *TILAM* is present in human HSCs by performing rapid amplification of cDNA ends (RACE) followed by RT-PCR to clone the full-length transcript. The cloned product has a structure similar to *ENSG00000249406* except for variations in internal splicing (Figure 1*A*, bottom, Supplementary Figure 1*B*). These results show that *TILAM* is expressed in HSCs, and there are multiple isoforms, which retain 5’ and 3’ conservation.

### TILAM is enriched in HSCs, induced by TGF-β signaling, and regulates COL1A1 expression

We found that *TILAM* was enriched in cultured primary human HSCs compared to normal liver and other primary tissues and cell lines (Figure 1*B*). *TILAM* is also induced by TGF-β signaling (Figure 1*B* and *C*), as described for *COL1A1*^40^. In addition, *TILAM* and *COL1A1* followed parallel expression patterns across tissues and cell types (Supplementary Figure 1*C*). *TILAM* is present in both the nucleus and cytoplasm, with more transcripts in the cytoplasm (Supplementary Figure 2*A* and *B*), and we designed locked nucleic acid antisense oligonucleotides (LNAs)^41^ to deplete transcripts in both compartments. Both LNAs successfully depleted *TILAM* in primary HSCs, and depletion was associated with reduced *COL1A1* mRNA (Figure 1*D*). We repeated depletion studies in primary human HSCs from three additional donors, which confirmed these findings (Supplementary Figure 2*C*-*E*). RNA-seq analysis revealed that *TILAM* depletion was associated with a broader reduction in expression of genes involved with ECM organization, cell adhesion, and motility, while genes involved in RNA processing/splicing were induced (Figure 1*E* and *F* and Supplementary Table 1). These results show that *TILAM* is induced by the profibrotic signal TGF-β, and depletion is associated with reduced expression of *COL1A1* and other ECM genes.

### lncRNA TILAM activity in liver organoid models

We next evaluated *TILAM* expression and activity in human liver organoids (HLOs), which include hepatocytes, HSCs, and cholangiocytes^38, 42^. We detected *TILAM* in HLOs under normal culture conditions and found that *TILAM* was induced with TGF-β treatment, along with *COL1A1* (Figure 2*A*). Using genome editing and homologous recombination, we created a human embryonic stem cell (hESC) line deficient in *TILAM* by inserting a cDNA encoding enhanced green fluorescent protein (GFP) followed by a polyadenylation (polyA) signal near the 5’ end of the gene to terminate transcription (Figure 2*B*). We applied this approach^43^ to allow transcription to continue at the lncRNA locus without production of the lncRNA product. *TILAM^gfp/gfp^*hESCs proliferated normally (data not shown) and were differentiated into HLOs. These HLOs also exhibited reduced expression of *COL1A1* in response to fibrotic signals (Figure 2*C*). We then depleted *TILAM* in HLOs differentiated from wild-type hESCs, which again led to reduced *COL1A1* expression (Figure 2*D*). These results provide further evidence that *TILAM* is regulated by TGF-β and promotes *COL1A1* expression in a 3D human organoid model.

**Figure 2.**
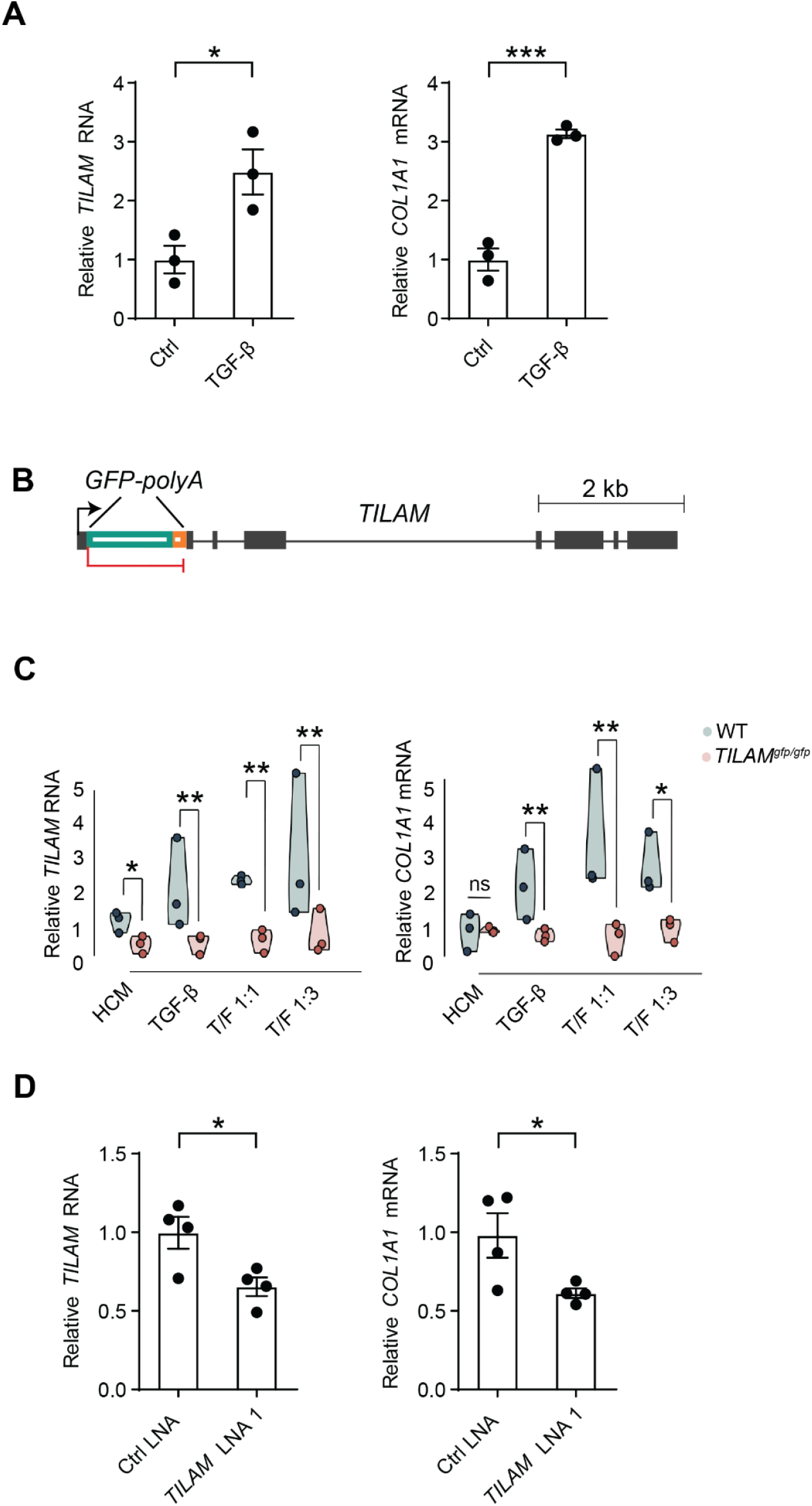
*TILAM* activity in human liver organoids. (*A*) Human embryonic stem cells (hESCs) were differentiated into liver organoids before treatment with TGF-β (10 ng/ml) for 4 days^38^. *TILAM* and *COL1A1* levels were quantified by qRT-PCR. *indicates p<0.05 and *** indicates p<0.001, (2-tailed unpaired *t* test). Error bars represent mean ± SEM. (*B*) Genome editing was performed to insert a cDNA encoding GFP and polyadenylation (polyA) signal into *TILAM* to disrupt expression of the lncRNA. (*C*) Expression of *TILAM* and *COL1A1* was quantified by qRT-PCR after wild-type (WT) and *TILAM*-deficient (*TILAM^gfp/gfp^*) hESCs were differentiated into liver organoids and cultured in the indicated conditions for 4 days compared to organoids remaining in hepatocyte culture media (HCM) alone. TGF-β 10 ng/ml (TGF); TGF-β 10 ng/ml and FGF-1 10 ng/ml (TGF/FGF 1:1); TGF-β 10 ng/ml and FGF-1 30 ng/ml (TGF:FGF 1:3). *ACTB* was used as an endogenous control. *indicates p<0.05, ** indicates p<0.01, (Kruskal-Wallis test with post-hoc Conover’s test). (*D*) hESCs were differentiated into liver organoids before delivery of *TILAM* LNA 1 or control LNA via gymnosis for 48 hours. Expression *of TILAM* and *COL1A1* was quantified by qRT-PCR. *indicates p<0.05, (2-tailed unpaired *t* test).

### Identification of a mouse ortholog of TILAM

There are currently no validated lncRNA transcripts annotated in a syntenic position to *TILAM* in the murine genome. To determine if there is an ortholog to *TILAM* near *Col1a1* in the murine genome, we performed RNA-seq and *ab initio* assembly of the transcriptome from HSCs isolated from mice before and after CCl_4_ treatment (Figure 3*A*). *Lrat-Cre* mice^8^ were crossed with *Zs-Green* Cre reporter mice to generate Zs-Green-labeled HSCs. HSCs were sorted from mice before and after four weeks of treatment with CCl_4_. RNA was analyzed by 100 nt paired-end RNA-seq. We assembled more than 8,500 genes in murine HSCs that did not overlap in sense with protein-coding genes, were greater than 200 nt in length, and had poor coding potential, consistent with lncRNAs (Figure 3*B* and Supplementary Table 2 and 3). These transcripts were also expressed at lower levels than mRNAs (Figure 3*C*, Supplementary Table 4).

**Figure 3.**
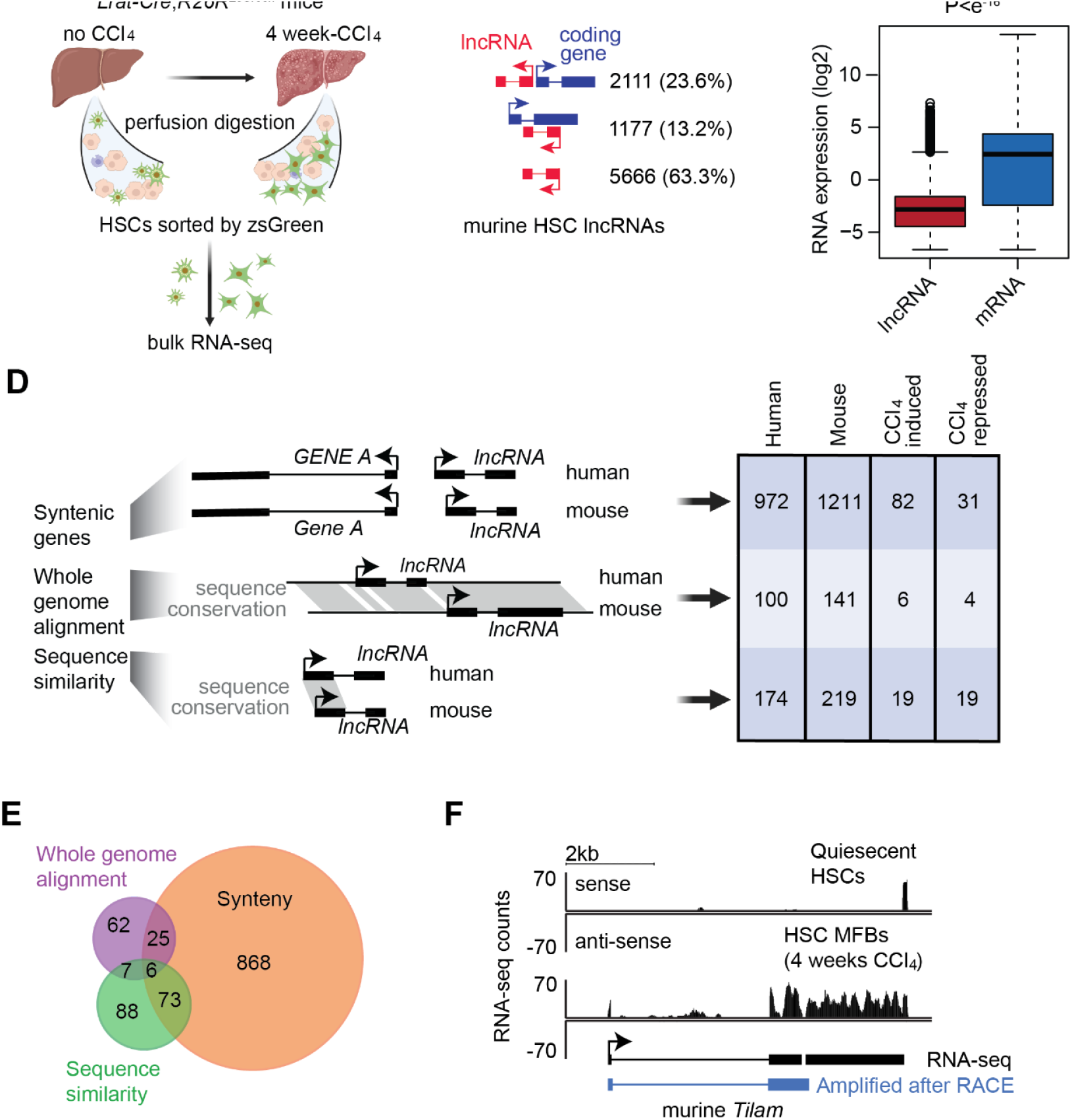
*TILAM* is conserved between human and murine HSCs. (*A*) Schematic showing the approach to generate RNA-seq data for HSCs *in vivo*. HSCs were sorted from mice (*Lrat-Cre;R26R^ZsGreen^*) before and after treatment with CCl_4_ for 4 weeks to induce liver fibrosis and differentiation of HSC myofibroblasts. Bulk RNA-seq data were generated from both healthy and fibrotic HSCs. (*B*) We identified 8954 putative lncRNAs in murine HSCs from RNA-seq data. lncRNAs are divided into divergent (top), natural antisense (center), and intergenic (bottom). (*C*) Expression levels of lncRNAs and mRNAs in murine HSCs. (*D*) We defined orthologous lncRNAs between human and murine HSCs by integrating three approaches: synteny (top), whole genome alignment (middle), and sequence similarity (bottom). The number of lncRNAs in each category are indicated as well as those induced or repressed with CCl_4_ treatment (right). (*E*) Venn diagram showing the number of human lncRNAs with murine orthologs as determined by the approaches in *D*. (*F*) RNA-seq tracks show reads mapped to *Tilam* in murine HSCs with or without 4 weeks of CCl_4_ treatment. RNA-seq counts are indicated on the y-axis, and assembled transcripts identified by RNA-seq (black) and amplified after RACE (blue) are shown below.

We identified orthologs through three approaches^44^: 1) genes in the same genomic position relative to protein-coding genes in human HSCs (synteny), 2) genes in regions of conserved genomic alignment, and 3) conserved exon sequence (sequence similarity) (Figure 3*D*, left). The first two categories do not rely on sequence similarity, and the majority of potential orthologs were identified through synteny (Figure 3*D*, right and 3*E*, Supplementary Figure 3*A* and Supplementary Table 5).

Through this analysis, we defined a spliced transcript that met criteria for an lncRNA adjacent to the *Col1a1* gene (Figure 3*F*). This gene was antisense to *Col1A1*, contained two exons, and was positioned ∼2.7 kb upstream of the *Col1a1* TSS (Supplementary Figure 3*B*). We confirmed that the annotated *Tilam* transcript exists in HSCs by RACE, followed by amplification of the full-length transcript from murine HSC RNA (Supplementary Figure 3*C*). RNA-seq annotated a second transcript downstream of the spliced lncRNA product (Figure 3*F*). The second transcript was separated from the 3’ end of the first transcript by a repetitive sequence that mapped to multiple areas of the genome (Supplementary Figure 3*D*), suggesting that the 63 nucleotide break between these two transcripts may have been created because sequencing reads did not align uniquely to this region. The product cloned by RACE bridged this gap, but we did not detect a transcript containing the full sequence of the two transcripts. The human and murine lncRNAs retained limited sequence conservation, with 117 nt of *TILAM* showing sequence conservation with 115 nt of *Tilam* (Supplementary Figure 3*E*).

Next, we quantified *Tilam* expression in bulk liver RNA isolated from mice receiving CCl_4_ compared to control treatment, which showed induction of *Tilam* with chronic injury (Figure 4*A*). Depletion of this lncRNA in murine HSCs also resulted in reduction of *Col1a1* gene expression (Figure 4*B*), similar to that observed with depletion of *TILAM* in human HSCs. Expression of *Tilam* was restricted to HSCs compared to 25 other cell types (Supplementary Figure 4*A* and *B*), with the exception of bladder tissue. These results identify an lncRNA expressed in murine HSCs with a similar genomic location, pattern of expression, and phenotype to human *TILAM*.

**Figure 4.**
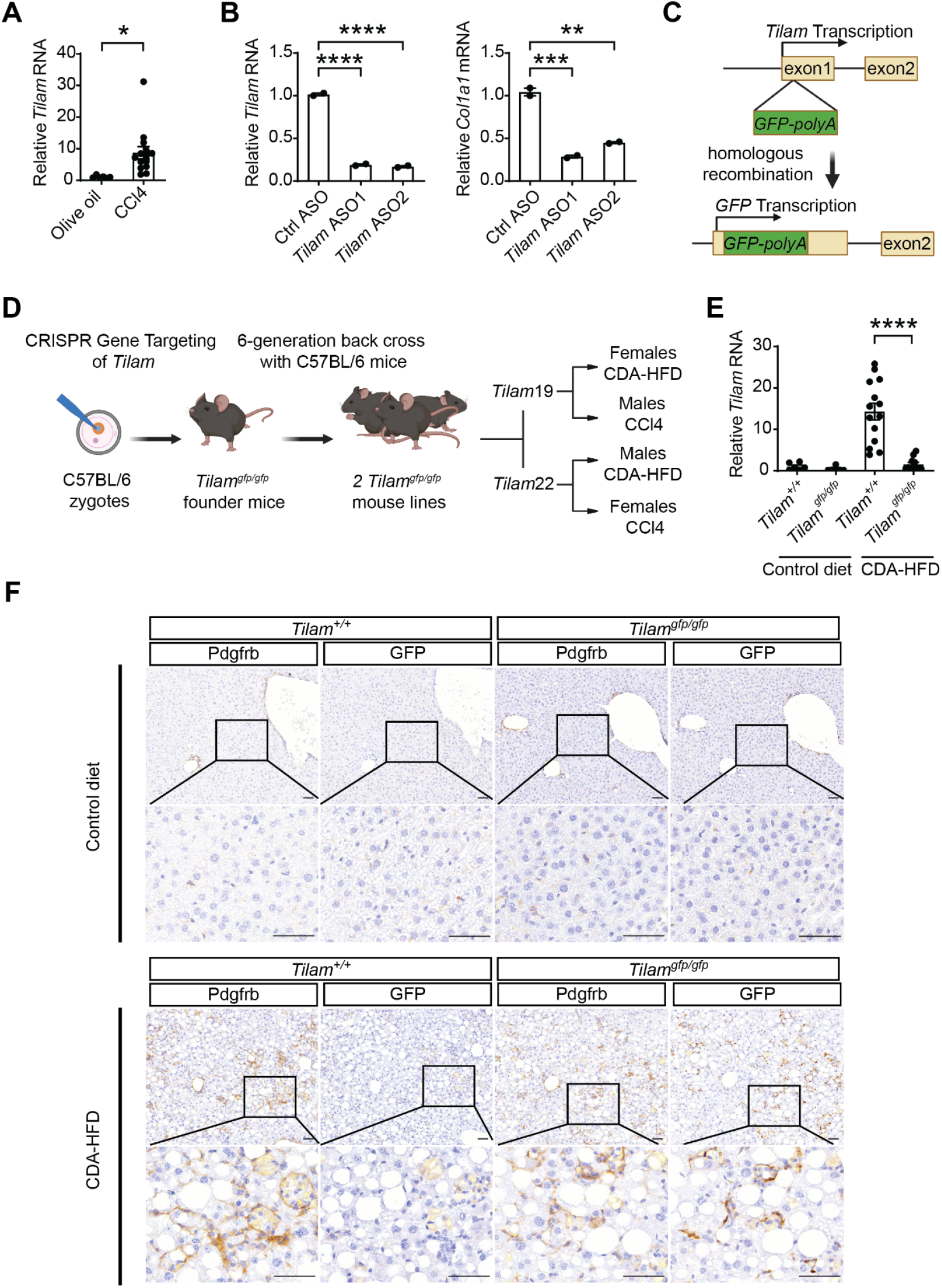
Generation of *Tilam^gfp/gfp^*mice. (*A*) qRT-PCR was performed on bulk liver from mice receiving CCl_4_ or control olive oil gavage for 4 weeks (n=6, 14). * indicates p <0.05, (2-tailed unpaired *t* test). Error bars represent mean ± SEM. (*B*) Murine HSCs were transfected with antisense oligonucleotides targeting *Tilam* (*ASO1*, *ASO2*) or non-targeting control ASO (*ctrl*). qRT-PCR was performed after 2 days to quantify *Tilam* and *Col1a1*. ** indicates p< 0.01, *** indicates p< 0.001, **** indicates p< 0.0001 (one-way ANOVA with Dunnett’s multiple comparison). (*C*) Targeted *GFP-polyA* replacement of the *Tilam* gene in mice. The *GFP-polyA* cassette was inserted between nt 55 and 56 to disrupt transcription of *Tilam*. (*D*) CRISPR targeting vector containing genomic regions flanking the *GFP-polyA* cassette were microinjected into the nucleus of C57BL/6 zygotes with guide RNAs and Cas9 protein. Two founder lines, *Tilam19* and *Tilam*22 were back crossed six generations and maintained on C57BL/6 background. Both genders of *Tilam^gfp/gfp^* mice were treated with CDA-HFD or CCl_4_ to induce liver fibrosis. (*E*) *Tilam* expression was quantified in *Tilam^+/+^* and *Tilam^gfp/gfp^* mice by qRT-PCR after 12 weeks of CDA-HFD treatment (n=6, 6, 14, 11). **** indicates p <0.0001, (2-tailed unpaired *t* test). (*F*) IHC was performed to detect Pdgfrb and GFP in adjacent murine liver sections in mice receiving CDA-HFD or control diet. Scale bar = 50 μm.

### In vivo mouse model to study Tilam

We created *Tilam-*deficient mice to determine if loss of *Tilam* could protect against the development of liver fibrosis *in vivo*. We applied genome editing to create founder lines in which a cDNA for GFP followed by a polyA signal (GFP-polyA) was inserted into the first exon of the *Tilam* gene to disrupt expression (Figure 4*C*). We confirmed three founder lines with heterozygous insertions in the correct genomic location (*Tilam^gfp/+^*) (Supplementary Figure 4*C, D*). Mice were generated on a C57BL/6 background and were backcrossed for six generations to wild-type C57BL/6 mice to reduce the chance that off-target effects from genome editing might affect the results. Heterozygous mice from the same founder were then intercrossed to generate *Tilac^gfp/gfp^* mice in which both alleles of *Tilam* were disrupted (Figure 4*D*).

The GFP-polyA sequence was inserted into the *Tilam* gene to disrupt expression of *Tilam*. To confirm this activity, we quantified *Tilam* expression in whole liver from *Tilam^+/+^* and *Tilam^gfp/gfp^* mice receiving CDA-HFD for 12 weeks to induce fibrosis (Figure 4*E*). GFP expression was then visualized by immunohistochemistry (IHC) (Figure 4*F*). Sections from livers of *Tilam^gfp/gfp^* (right) and *Tilam^+/+^* mice (left) receiving either control diet or CDA-HFD were stained for PDGFRβ to mark HSCs and GFP to mark cells that would normally express *Tilam.* Staining in serial sections revealed induction of GFP in areas of PDGFRβ expression in *Tilam^gfp/gfp^* mice receiving CDA-HFD but no GFP expression in wild-type mice with CDA-HFD. *Tilam^gfp/gfp^* mice receiving normal diet showed only trace expression of GFP near vessels (Figure 4*F* and Supplementary Figure 5*A*). These results show that *Tilam* is induced primarily in HSCs with fibrotic liver injury, and *Tilam* expression is significantly reduced in *Tilam^gfp/gfp^* mice.

### Tilam deficiency results in attenuated liver fibrosis in vivo

*Tilam^gfp/gfp^* mice were next challenged with CDA-HFD to determine if depletion of *Tilam* protects against the development of liver fibrosis. We observed a significant reduction in hydroxyproline levels in the livers of *Tilam^gfp/gfp^* compared to wild-type controls with hydroxyproline levels decreasing from 168 μg/g (standard error of the mean, SEM 8.8) in control mice to 120 μg/g

(SEM 8.4) in *Tilam^gfp/gfp^* mice (Figure 5*A*). Analysis of collagen proportionate area (CPA) based on Sirius red staining also showed reduced fibrosis in *Tilam^gfp/gfp^* mice (Figure 5*B*) with a CPA score of 5.4% (SEM 0.31) for control and 3.6% (SEM 0.42) for *Tilam^gfp/gfp^* mice. Representative histologic sections show increased Sirius red staining in control mice receiving CDA-HFD compared to control mice with normal diet. *Tilam^gfp/gfp^*mice receiving CDA-HFD showed increased Sirius red compared to normal diet, but this staining is reduced compared to wild-type control mice receiving CDA-HFD (Figure 5*C*). Gene expression from bulk liver also demonstrated reduced expression of *Col1a1*, *Acta2*, and *Timp1* (Figure 5*D*-*F*), while *Tgfb1* and *Mmp2* levels were not significantly changed (Supplementary Figure 5*B* and *C*).

**Figure 5.**
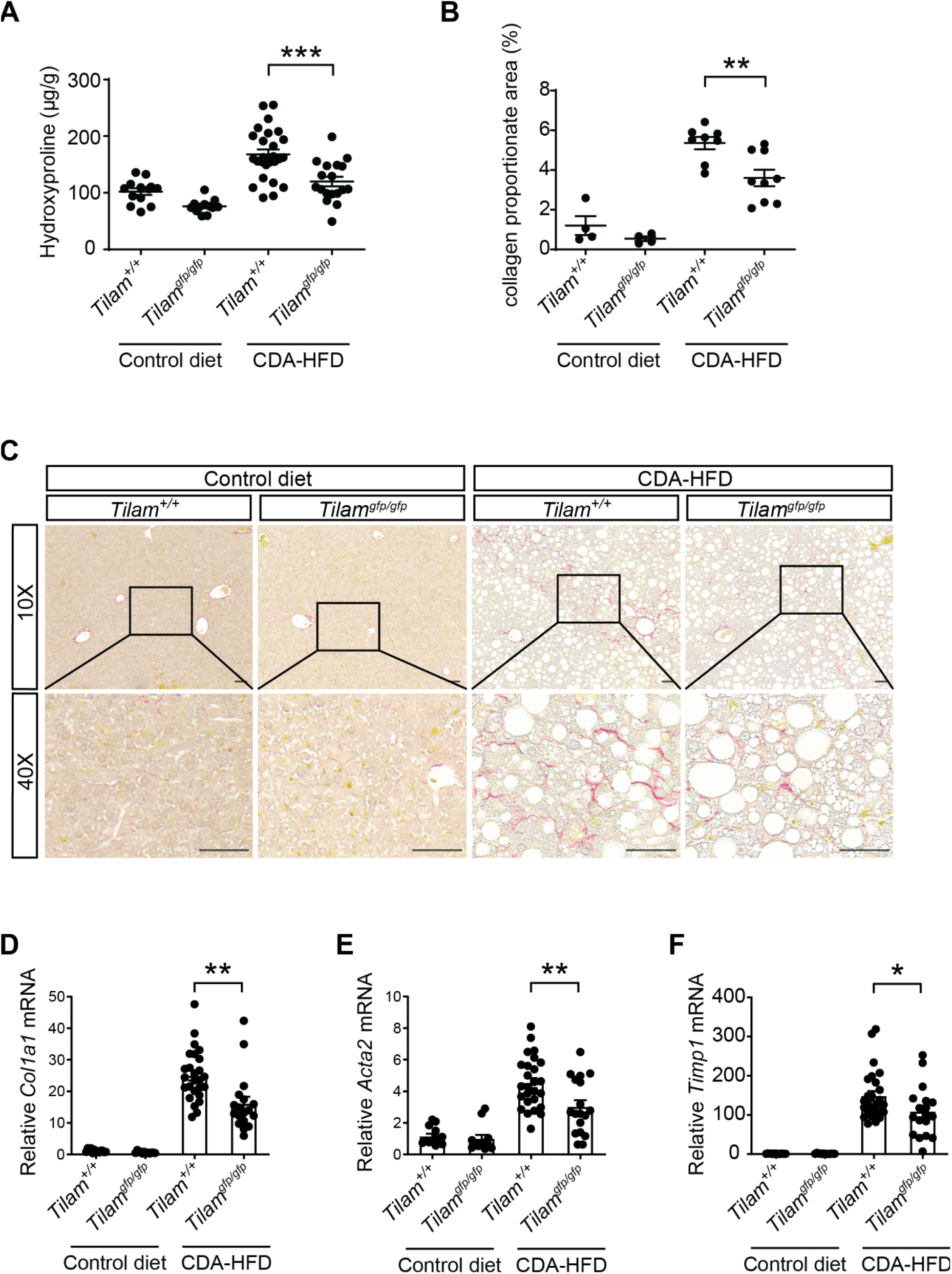
Loss of *Tilam* protects against liver fibrosis in CDA-HFD female mice. (*A*) Hydroxyproline was quantified in *Tilam^gfp/gfp^* and *Tilam^+/+^* female mice (*Tilam19*) after 12 weeks of CDA-HFD or control diet, *Tilam^gfp/gfp^* livers showed a significant reduction in hydroxyproline levels, compared to *Tilam^+/+^*(n=13, 12, 26, 18). *** indicates p<0.001 (2-tailed unpaired *t* test). Error bars represent mean ± SEM. (*B*) Collagen proportionate area (CPA) was calculated for the indicated conditions. Samples closest to the mean for each condition in (A) were selected for analysis (n=4, 4, 8, 9). ** indicates p<0.01 (2-tailed unpaired *t* test). (*C*) Representative images of Sirius red staining in *Tilam^gfp/gfp^* and *Tilam^+/+^* female mice with control and CDA-HFD. Scale bar = 50 μm. (*D*-*F*) *Col1A1*, *Acta2,* and *Timp1* expression for bulk liver were quantified by qRT-PCR from the CDA-HFD-treated female cohort (n=13, 12, 26, 18). *indicates p<0.05, ** indicates p<0.01 (2-tailed unpaired *t* test).

These studies were performed using female mice from founder line one (*Tilam19*). Male *Tilam^gfp/gfp^* mice from founder line one also showed reduced hydroxyproline (Figure 6*A*) and *Col1A1* mRNA (Figure 6*B*) levels in the CCl_4_ fibrosis model. We repeated the CDA-HFD experiment with male mice from founder line two (*Tilam22*) (Figure 6*C*) and the CCl_4_ experiment with female mice from founder line two (Figure 6*D*). In all experiments, *Tilam^gfp/gfp^*mice demonstrated reduced hydroxyproline levels compared to controls. Together, *in vivo* studies showed that reduction in the *Tilam* transcript protects against the development of fibrosis even when the locus remains actively transcribed.

**Figure 6.**
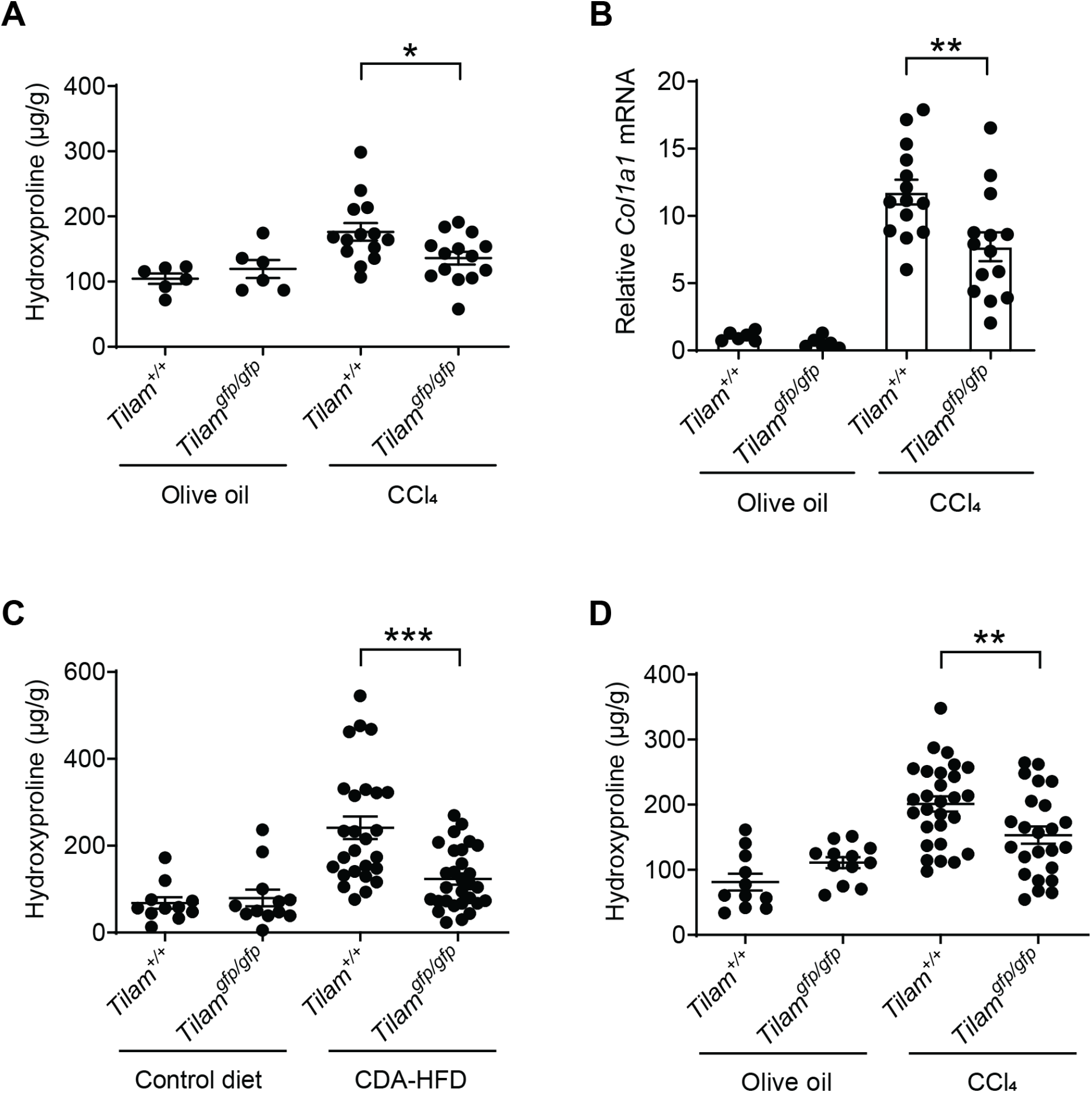
Loss of *TILAM* protects against liver fibrosis in other mouse groups with CCl_4_ and CDA-HFD treatment. (*A-B*) Hydroxyproline and *Col1a1* expression levels in male mice (*Tilam19*) after 4 weeks of CCl_4_ treatment (n=6, 6, 14, 14). *indicates p<0.05, ** indicates p<0.01 (2-tailed unpaired *t* test). Error bars represent mean ± SEM. (*C*) Hydroxyproline analysis in male *Tilam^+/+^*and *Tilam^gfp/gfp^* livers (*Tilam22*) after 12 week CDA-HFD treatment (n=11, 12, 26, 29). *** indicates p<0.001, (2-tailed unpaired *t* test). (*D*) Hydroxyproline levels in female *Tilam^+/+^*and *Tilam^gfp/gfp^* livers (*Tilam22*) after 4 weeks of CCl_4_ treatment (n=11, 12, 28, 24). ** indicates p<0.01 (2-tailed unpaired *t* test).

### TILAM interacts with PML to regulate fibrosis

To define interacting partners that could explain the activity of *TILAM*, we expressed *TILAM* fused to a tandem S1m aptamer (*TILAM-apt*) recognizing streptavidin^21^ and a scrambled sequence of *TILAM* fused to the same aptamer (*scTILAM-apt,* Supplementary Figure 6) in LX-2 stellate cells using lentivirus. Mass spectrometry (MS) was performed to identify the proteins enriched in precipitates from *TILAM-apt* compared to *scTILAM-apt* (Figures 7*A* and 7*B* and Supplementary Table 6). We then depleted mRNAs encoding the proteins most enriched in the MS analysis (Figure 7*B*, circled) to determine which gene products affected expression of *COL1A1* (Figure 7*C*). This analysis revealed that depletion of PML was associated with the greatest reduction in *COL1A1*. These results suggest that *TILAM* interacts with PML to regulate the fibrotic activity of HSCs and prompted further analysis.

**Figure 7.**
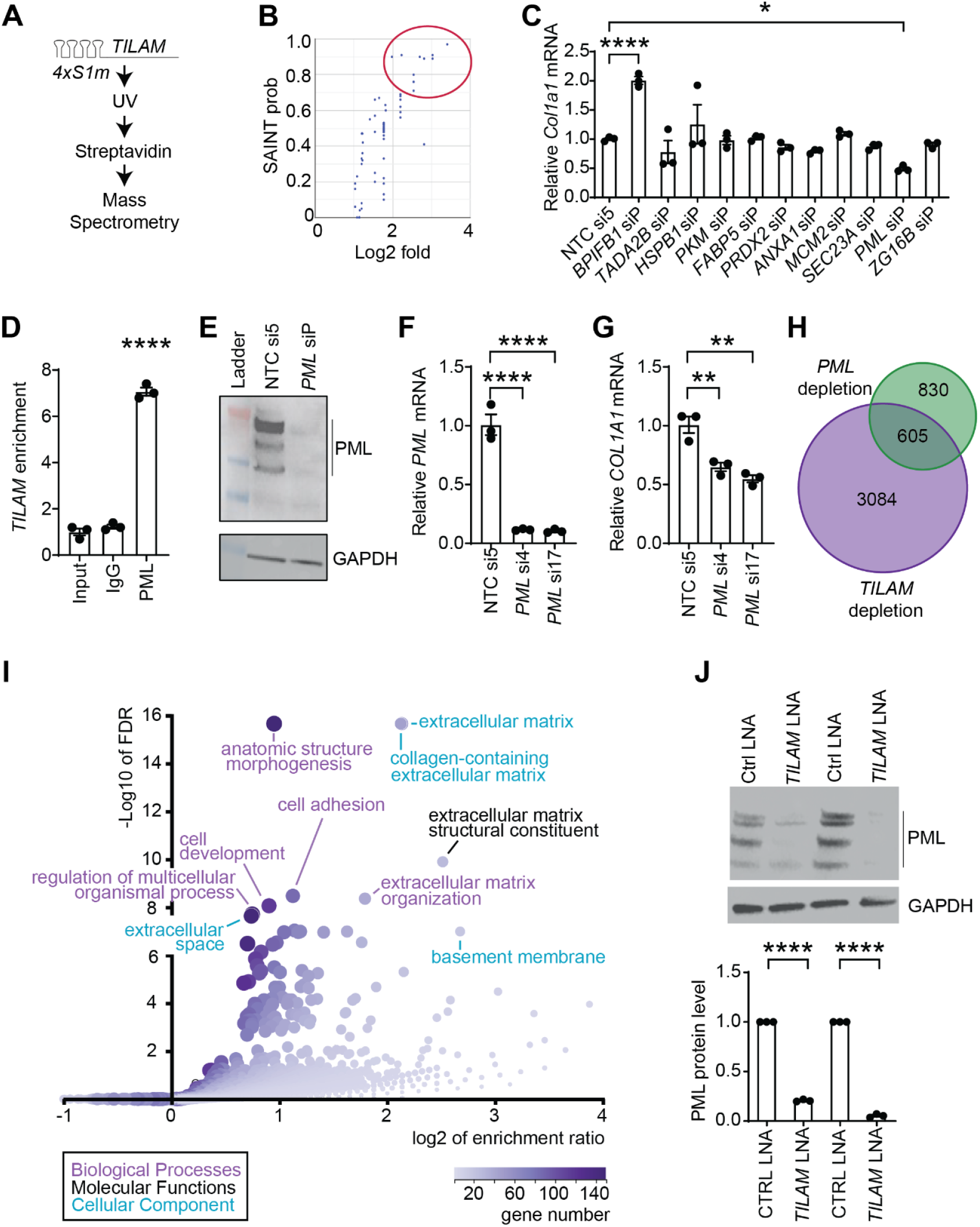
*TILAM* interacts with PML to regulate the fibrotic activity of HSCs. (*A*) Schematic showing approach to identify proteins that interact with *TILAM* using the 4xS1m aptamer (*TILAM-apt*). (*B*) MS analysis for *TILAC-apt* compared to a scrambled (sc) *TILAM-apt* in LX-2 cells. The probability score represents the significance analysis of interactome (SAINT) score (y-axis) and is displayed versus the log 2 fold enrichment (x-axis). The red circle indicates the proteins most enriched with precipitation of *TILAM-apt.* (*C*) mRNAs encoding the proteins most enriched with *TILAM-apt* precipitation were depleted in primary human HSCs using pooled siRNAs, and *COL1A1* expression was quantified by qRT-PCR. * indicates p<0.05, *** indicates p<0.001 (one-way ANOVA with Dunnett’s multiple comparison). Error bars represent mean ± SEM. (*D*) RNA immunoprecipitation was performed in primary human HSCs using an antibody against PML and IgG control. *TILAM* RNA was quantified with each precipitation and compared to cell lysates (input). **** indicates p<0.0001 (one-way ANOVA). (*E*) PML protein was quantified by Western blot following depletion with *PML* siP compared to non-targeting control (si5). The molecular weight ladder is included (left). GAPDH was evaluated as a loading control. (*F*) *PML* mRNA was quantified by qRT-PCR following transfection of primary human HSCs with two different siRNAs targeting *PML* (si4, si17) and compared to a non-targeting control (si5). **** indicates p<0.0001 (one-way ANOVA). (*G*) *COL1A1* mRNA is quantified under the same conditions as *E*. ** indicates p<0.01 (one-way ANOVA). (*H*) RNA-seq was performed on primary human HSCs depleted of either *TILAM*, *PML*, or treated with respective control oligos. Venn diagram shows the number of genes that decrease in expression with depletion of *TILAM* (purple) and *PML* (green). The intersection indicates genes in common between depletion of *TILAM* and *PML*. (*I*) GO analysis was performed on the 605 genes repressed with depletion of both *TILAM* and *PML*. Each circle represents a GO category, which is plotted to indicate −log10FDR (y-axis) versus log2 fold enrichment (x-axis). The color of each dot represents the number of genes in a GO category, and the size indicates the number of genes affected in a category. Individual categories are highlighted, and the color of the label indicates the source of the category (lower left). (*J*) PML protein was visualized by Western blot in primary human HSCs following depletion of *TILAM* (top). Biologic replicates are shown. GAPDH was probed as a loading control. Band intensities were quantified (bottom and normalized to GAPDH signal). Each band was quantified three times. **** indicates p<0.0001, (2-tailed unpaired *t* test).

Precipitation of PML protein in primary HSCs followed by qRT-PCR confirmed that endogenous *TILAM* interacts with endogenous PML (Figure 7*D*). Depletion of PML with pooled siRNAs in HSCs from a second donor was also associated with reduced *COL1A1* expression (Supplementary Figure 7*A* and *B*). We confirmed that depletion of *PML* mRNA reduced PML protein (Figure 7*E*) and then depleted *PML* using two individual siRNA duplexes to provide further confirmation that depletion of PML was associated with reduced *COL1A1* expression (Figure 7*F*-*G*). These results show that endogenous *TILAM* interacts with PML in primary human HSCs, and depletion of PML leads to reduced expression of *COL1A1* in multiple primary HSCs lines.

To understand the overlap in gene pathways downstream of *TILAM* and *PML*, we depleted *TILAM* (control LNA versus LNA 1) and *PML* (control siRNA versus siRNA 17) separately and performed RNA-seq. We identified 605 genes that were repressed in common with depletion of *TILAM* and *PML* (Figure 7*H*). Gene Ontology (GO) revealed that both *TILAM* and PML regulate pathways involved in ECM production and cell adhesion (Figure 7*I* and Supplementary Figure 7*C*). Depletion of PML alone, which is found predominantly in the nucleus of HSCs, also demonstrated enrichment in pathways regulating ECM production and cell adhesion (Supplementary Figure 7*D* and *E*). These results suggest that the interaction between *PML* and *TILAM* promotes ECM production.

### TILAM regulates PML protein levels

RNA-seq analysis revealed that depletion of *TILAM* was associated with an approximately two-fold reduction in *PML* mRNA (Supplementary Table 1). We next asked how depletion of *TILAM* affected PML protein levels. While *PML* mRNA levels were reduced by half, 85% or more of PML protein was lost with depletion of *TILAM* (Figure 7*J*), suggesting that *TILAM* may regulate ECM production in HSCs at least in part by maintaining PML protein levels.

## Discussion

Fibrosis is the strongest predictor of morbidity and mortality in chronic liver disease^45, 46^. Moreover, complications that arise with progression to cirrhosis lead to a reduced quality of life and increased economic burdens for patients and society^47^. While treatment of the underlying cause of disease can inhibit progression of fibrosis, and in some cases lead to regression, there remain sources of chronic disease with limited or unavailable treatments. Regardless of the source of injury, inhibiting the development or promoting the regression of fibrosis has the potential to reduce the morbidity and mortality associated with chronic liver disease.

HSCs are the primary cell type responsible for hepatic fibrosis^6–8^. We previously mapped lncRNAs expressed in human HSCs to identify candidate lncRNAs that may regulate fibrotic activity^39^. Here we focused on defining the function of *TILAM*, which we identified by its expression in HSC myofibroblasts, co-expression with *COL1A1,* and induction by TGF-β signaling^39^. Proximity and co-expression with *COL1A1* suggested that *TILAM* may regulate *COL1A1*, and we evaluated this possibility with two separate approaches in human HSCs. We first depleted *TILAM* using LNAs targeting the 3’ region of the transcript to avoid potential effects on *TILAM* transcription^48^. We then applied genome editing to insert a cDNA encoding GFP and a polyA signal^43^ to prevent production of *TILAM* while maintaining gene expression at the *TILAM* locus. Both approaches showed that depletion of *TILAM* leads to a decrease in *COL1A1*, indicating that the *TILAM* transcript is required to promote *COL1A1* expression. Furthermore, the effect of *TILAM* depletion on *COL1A1* expression was observed in multiple primary human HSC lines and in HLOs.

Only about a third of lncRNAs are conserved between humans and mice^49^. We performed RNA-seq and *ab initio* assembly of the murine transcriptome in quiescent HSCs and HSC myofibroblasts isolated *in vivo* to determine if an lncRNA could be identified that was induced in HSC myofibroblasts near *Col1a1*. This analysis defined a gene composed of two exons that is divergent to *Col1a1* but located ∼2.7 kb from *Col1a1* in the murine genome compared to ∼7.3 kb in the human genome. Poor sequence conservation between the mouse and human orthologs is frequently observed (Supplementary Figure 3*E*)^50^, but depletion of each transcript was associated with reduced *COL1A1* in their respective HSCs, leading to the conclusion that they are functional orthologs.

*Tilam* was disrupted in *Tilam^gfp/gfp^* mice by inserting a cDNA encoding GFP and a polyA signal into the first exon of *Tilam*. These mice developed and bred normally in the absence of *Tilam*, which was anticipated due to very limited expression detected in RNA-seq data from mouse tissue. In addition to preventing expression of *Tilam*, the insertion of GFP under the *Tilam* promoter allowed GFP to act as a reporter for *Tilam* expression and confirmed that *Tilam* is restricted to HSC myofibroblasts in the liver. *Tilam* expression was also noted in normal mouse bladder tissue based on RNA-seq (Supplementary Figure 4*A*), and additional studies will be required to confirm these findings and define the role of *Tilam* in the bladder. Future investigation will also be required to understand if *Tilam* expression and its ability to promote ECM production are restricted to the liver or are features of fibroblasts in other organs.

*COL1A1* is the nearest gene to *TILAM,* but it is not the only gene regulated by *TILAM*. Depletion of *TILAM* also reduces expression of other ECM genes including *COL1A2, COL3A1, COL4A2, COL5A1, TGFB2,* and *FN1,* suggesting that *TILAM* functions through a mechanism that reaches many different genes. Through MS studies, PML was identified as an interacting partner that could explain these observations. Initial studies were performed in LX-2 cells with ectopic expression of *TILAM*, and a scrambled version of *TILAM* was selected as a control to limit noise that might be created by proteins that bind RNAs of similar size or similar nucleotide content. Ectopic expression of *TILAM* in this analysis might also lead to higher levels of the lncRNA than normally present endogenously, which could facilitate binding events that do not occur in primary HSCs. Enrichment of *TILAM* with precipitation of PML in primary human HSCs confirms that endogenous PML interacts with endogenous *TILAM*.

PML is found in both the nucleus and cytoplasm^51^. In the nucleus, PML forms nuclear bodies (NBs), which can control gene expression^52^, and we observe the most intense signal for PML in NBs of human HSCs (Supplementary Figure 7*E*). The change in ECM mRNA expression observed with depletion of PML is consistent with regulation at the level of transcription or RNA stability, and the interaction between nuclear PML and Pin1 in cardiac fibroblasts provides an example of how PML can regulate RNA levels to promote fibrosis in a different system^53^. The current findings also cannot exclude the possibility that cytoplasmic interactions, such as those promoting TGF-β signaling^54^ could provide an alternative path for PML complexes to regulate gene expression less directly.

*TILAM* is necessary to maintain PML protein levels in HSC myofibroblasts. PML turnover is regulated by multiple mechanisms including phosphorylation and SUMOylation, which promote ubiquitination and degradation^55, 56^, and lncRNAs such as *HOTAIR* can also directly interact with E3 ubiquitin ligases^57^. PML can also be regulated at the level of translation through mediators including K-RAS and mTOR^58^. It is not yet clear how *TILAM* regulates PML protein levels and whether the interaction between *TILAM* and PML functions to promote expression of ECM genes or the interaction is only required to maintain PML protein, which regulates gene expression.

In summary, we have focused on identifying lncRNAs that promote the fibrotic activity of HSC myofibroblasts, as these have potential as therapeutic targets to inhibit progression of fibrosis through ASO technology such as siRNAs^59, 60^ or LNAs^61^. Retinoid-conjugated nanoformulations have also improved delivery of ASOs to HSCs^62, 63^, and targeting RNAs such as *TILAM*, which are restricted in expression to HSC myofibroblasts can provide cell-type-specific activity while reducing the risk of off target effects.

## Acknowledgments

The authors thank Bryan Fuchs, Disha Badlani, and Amin Mahpour for helpful discussions. We thank Kathrin Leppek and Georg Stoecklin for sharing the 4xS1m plasmid. We thank Mantu Bhaumik, Yiping Zhou, and the IDDRC Gene Manipulation & Genome Editing Core (NIH-P50 HD105351) for generation of *Tilam^gfp/gfp^* mice. We thank Ross Tomaino and the staff at the Taplin Mass Spectrometry Core at Harvard Medical School for performing MS analysis. Sirius red staining and IHC for mouse tissue was performed by iHisto. RNA-seq was performed through the Massachusetts General Hospital (MGH) Next Generation Sequencing Core and GENEWIZ. A.C.M. and Y.V.P. were supported by NIH/NIDDK grant R01DK116999. Y.V.P. was also supported by a grant from PSC Partners Seeking a Cure Canada. A.C.M. was also supported by a Pew Biomedical Scholars Award, DK043351 P30 Pilot Award, MGH Transformative Scholars Award, and internal MGH funding. R.F.S. was supported by the Columbia University Digestive and Liver Disease Research Center grant 5P30DK132710 and 5R01DKL12895. Figure 3*A*, Figure 4*C*, 4*D*, Supplementary Figure 4*C*, and graphical abstract were created with BioRender.com.

## Datasets

The GEO accession number for RNA-seq from human and murine HSCs performed for this paper will be GSE238159 upon release. Previously published datasets of human tissue and cells were obtained from dbGAP (GTEx)^64^, GSE26284, and GSE41009. Mouse tissue datasets were obtained from GSE36025 and GSE36114.

---

**Supplementary Figure 1.**
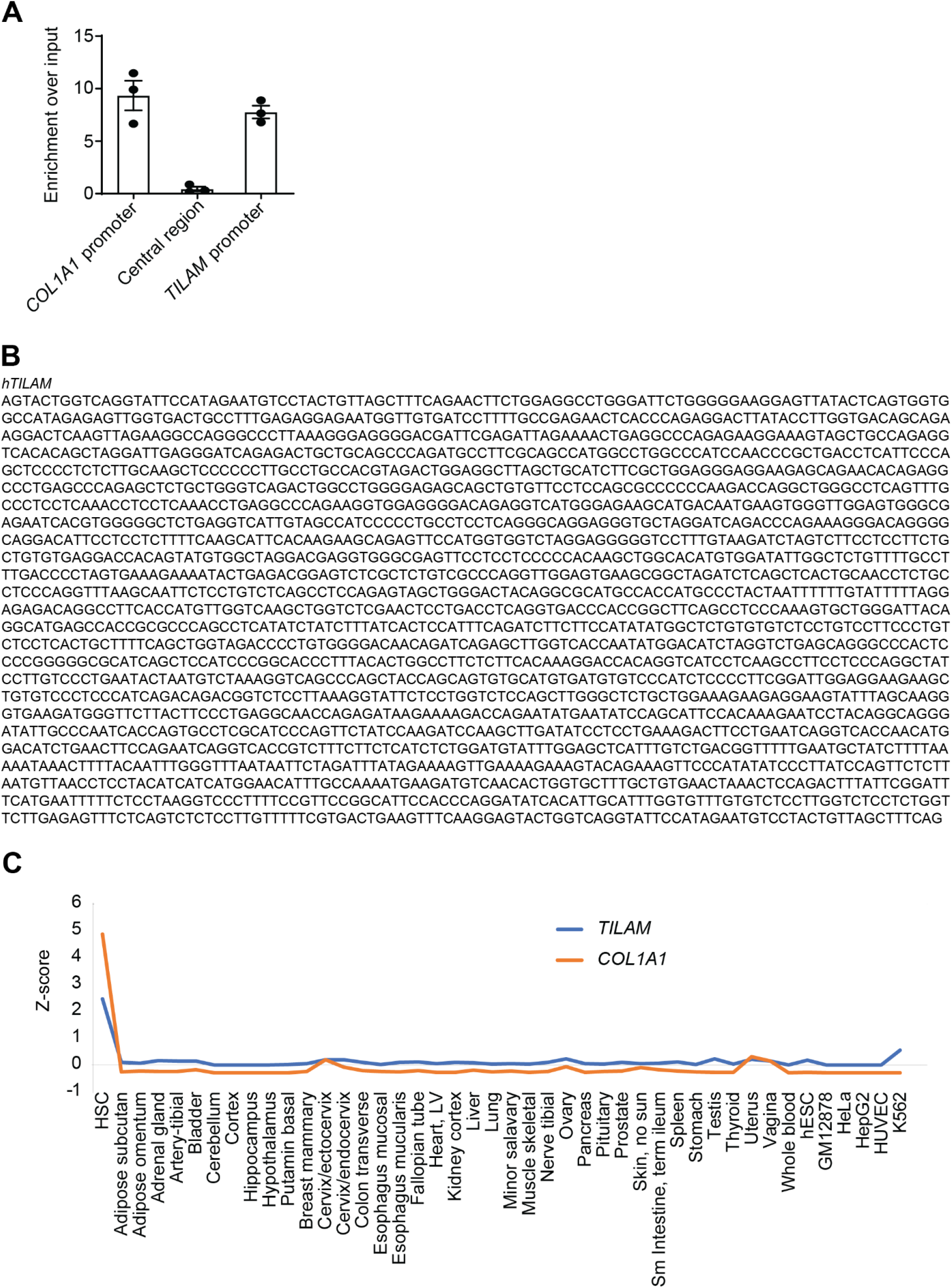
Accessibility between *COL1A1* and *TILAM*, *TILAM* sequence cloned from human HSCs, and expression across tissues and cell types. (*A*) Formaldehyde assisted isolation of regulatory elements (FAIRE)^1^ was performed followed by genomic qPCR to quantify relative accessibility (y-axis) of the *COL1A1* promoter, the *TILAM* promoter, and the region between both promoters (central region). Error bars represent mean ± SEM. (*B*) Sequence of the isoform of *TILAM* amplified from primary human HSCs. (*C*) *TILAM* and *COL1A1* expression across tissues and cell types (GTEx^2^, GSE26284, and GSE41009)^3^.

**Supplementary Figure 2.**
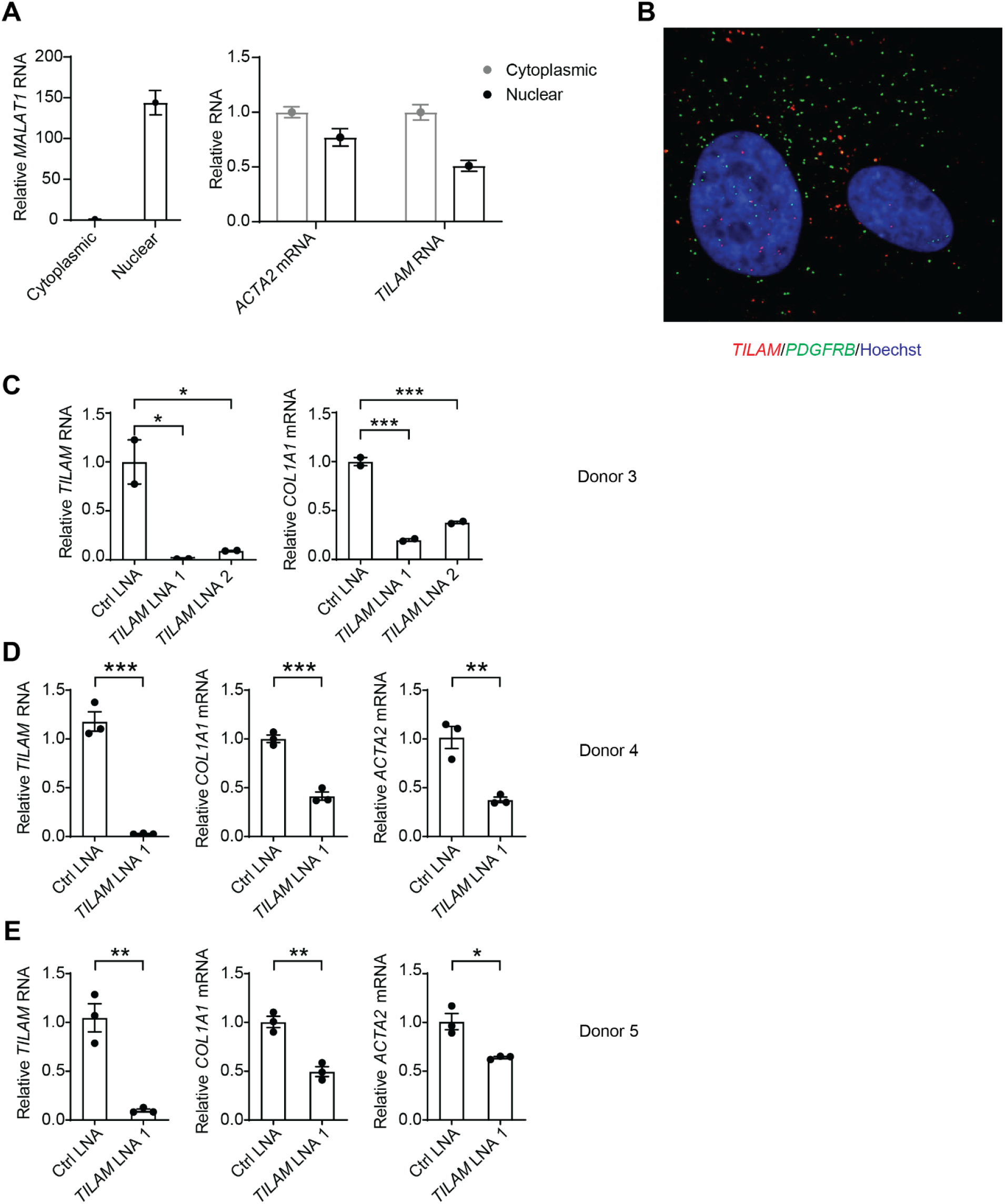
Subcellular localization of *TILAM,* and depletion of *TILAM* in multiple primary human HSC lines. (*A*) Nuclear and cytoplasmic fractions from primary human HSCs were separated, and *MALAT1* transcripts were quantified by qRT-PCR (left) to assess efficiency of fractionation, as *MALAT1* is retained in the nucleus. The distribution of *ACTA2* and *TILAM* RNA transcripts were quantified in the nuclear and cytoplasmic fractions by qRT-PCR (right). Error bars represent mean ± SEM. (*B*) Single molecule RNA FISH (smFISH) was performed using probes that recognize *TILAM* (red) and *PDGFRB* (green). Nuclei are stained with Hoechst (blue). (*C*) Primary human HSCs from donor 3 were transfected with LNAs targeting *TILAM* (LNA 1 and LNA 2) and a control LNA before *TILAM* and *COL1A1* were quantified by qRT-PCR. * indicates p<0.05 and *** indicates p<0.001 (one-way ANOVA with Dunnett’s multiple comparisons test). (*D*-*E*) Primary human HSCs from two additional donors were transfected with *TILAM* LNA 1 and control LNA before *TILAM*, *ACTA2*, and *COL1A1* were quantified by qRT-PCR. * indicates p<0.05, ** indicates p<0.01, *** indicates p<0.001 (2-tailed unpaired *t* test).

**Supplementary Figure 3.**
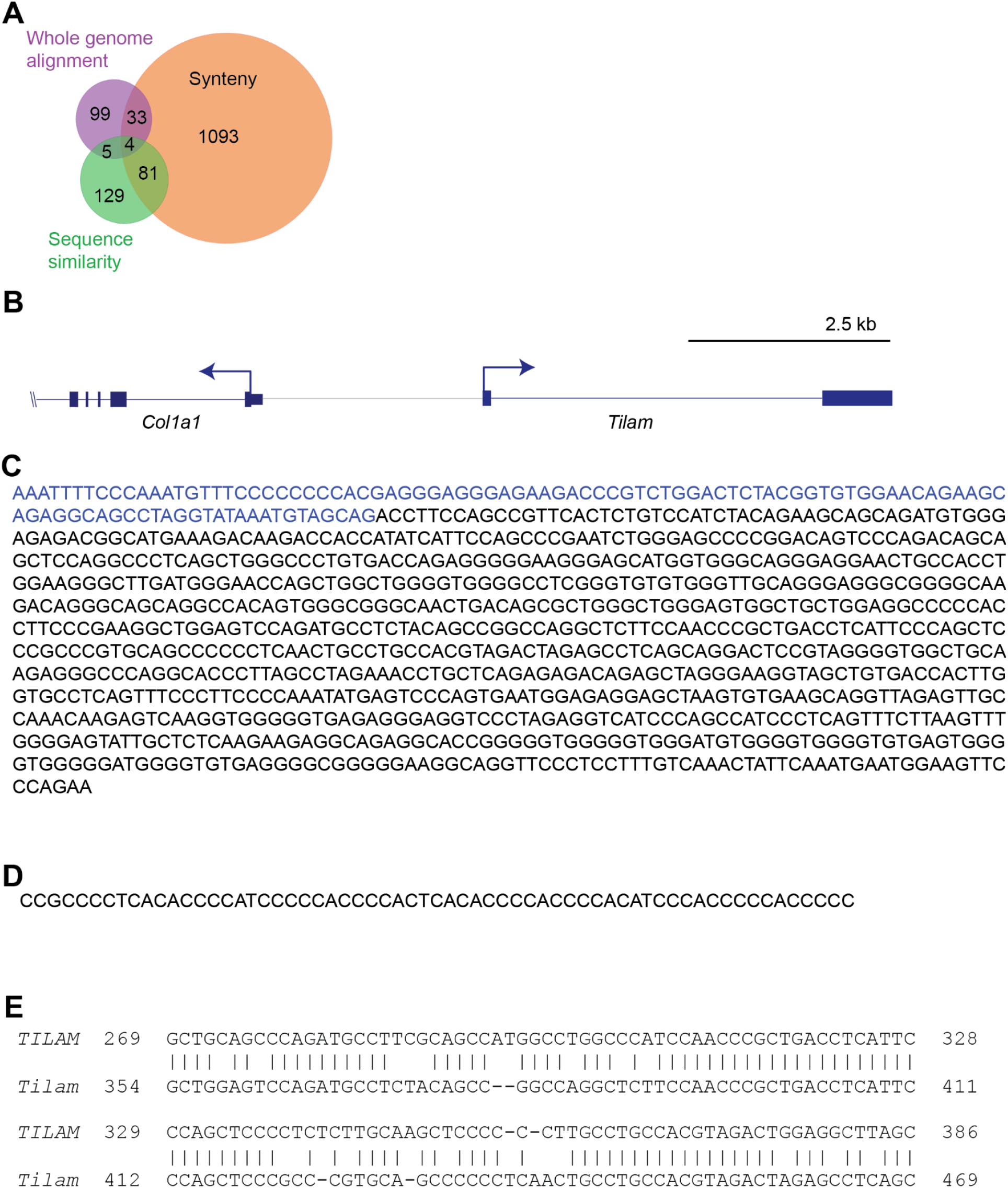
Sequence of *Tilam* and alignment with *TILAM*. (*A*) Venn diagram showing murine lncRNAs with human orthologs as determined by synteny, whole genome alignment, and sequence similarity. (*B*) Schematic of *Tilam* in relation to *Col1a1* in the mouse genome. (*C*) Sequence of *Tilam* cloned from murine HSCs. (*D*) Repetitive sequence in 63 nt gap between two transcripts in the *Tilam* locus assembled with RNA-seq data from murine HSCs (Figure 3F). (*E*) Sequence conservation between human (top) and murine *(bottom) TILAM* at indicated nucleotide positions. Nucleotide positions are based on cloned sequences (Supplemental Figure 1B and Supplemental Figure 3C).

**Supplementary Figure 4.**
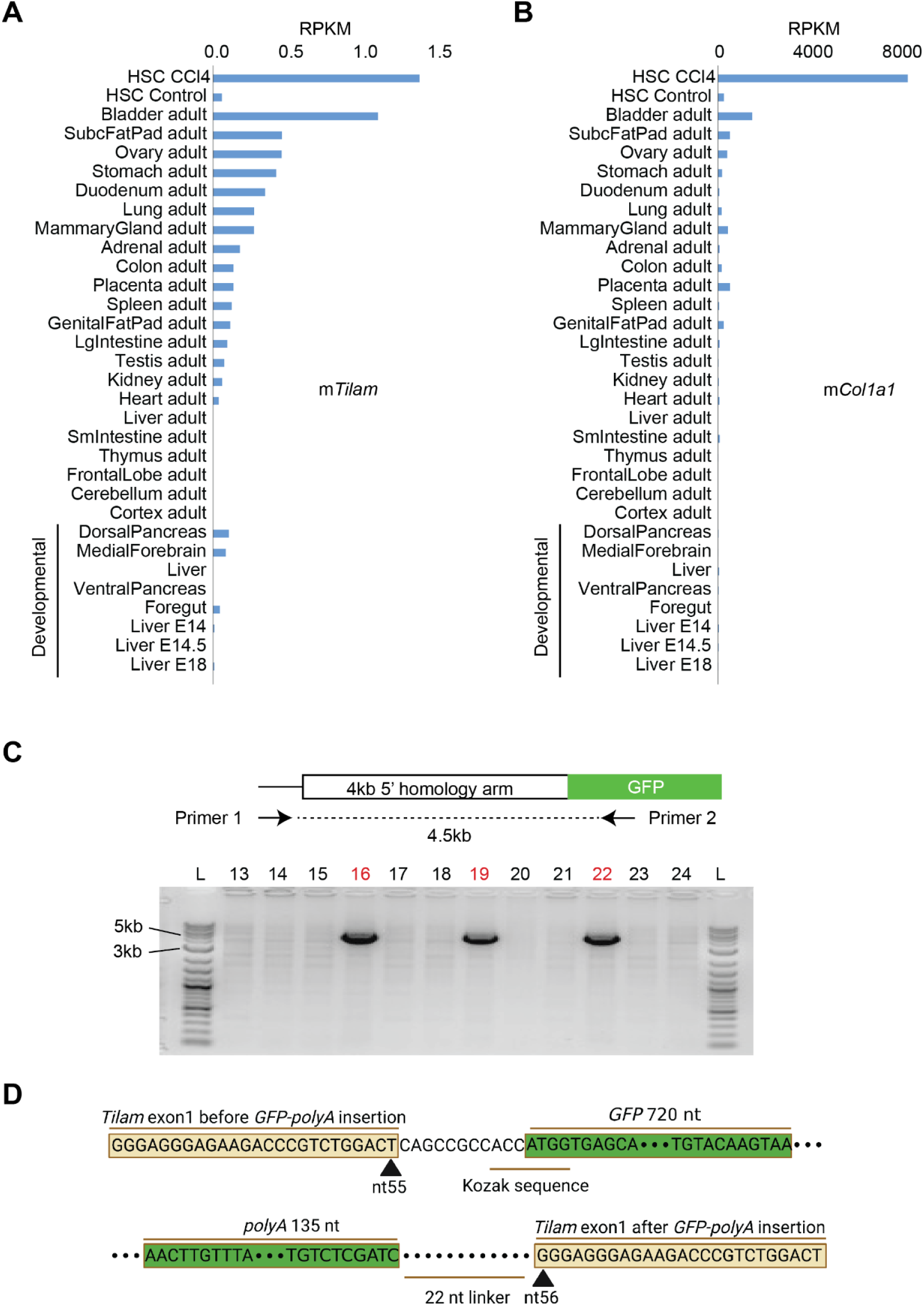
*Tilam* expression and insertion of *GFP* and a polyadenylation (polyA) signal to disrupt the *Tilam* gene. (*A*-*B*) Murine *Tilam* and *Col1a1* expression patterns across adult and developmental tissues (GSE36025 and GSE36114) compared to murine HSCs in this study. (*C*) Genomic PCR performed on pups born after zygote injections for genome editing to identify potential founders. The upstream primer (Primer 1) annealed to a region located outside the 4 kb homology arm used for homologous recombination while the downstream primer (Primer 2) annealed to a region within the *GFP* sequence. Products were visualized on an agarose gel. Mice 16, 19, and 22 (red) showed evidence of the correct insertion. These mice were crossed to wild-type C57BL/6 mice to confirm germline transmission and back crossed for six generations to wild-type C57BL/6 mice before intercrossing mice from the same founders. DNA ladder (L) is on the left and right, with 3kb and 5kb markers indicated on the left. (D) Sequencing of genomic DNA was performed in *Tilam^gfp/gfp^* mice, showing insertion of *GFP*-polyA sequence at nt 55 of exon 1. The sequence at indicated junctions is shown.

**Supplementary Figure 5.**
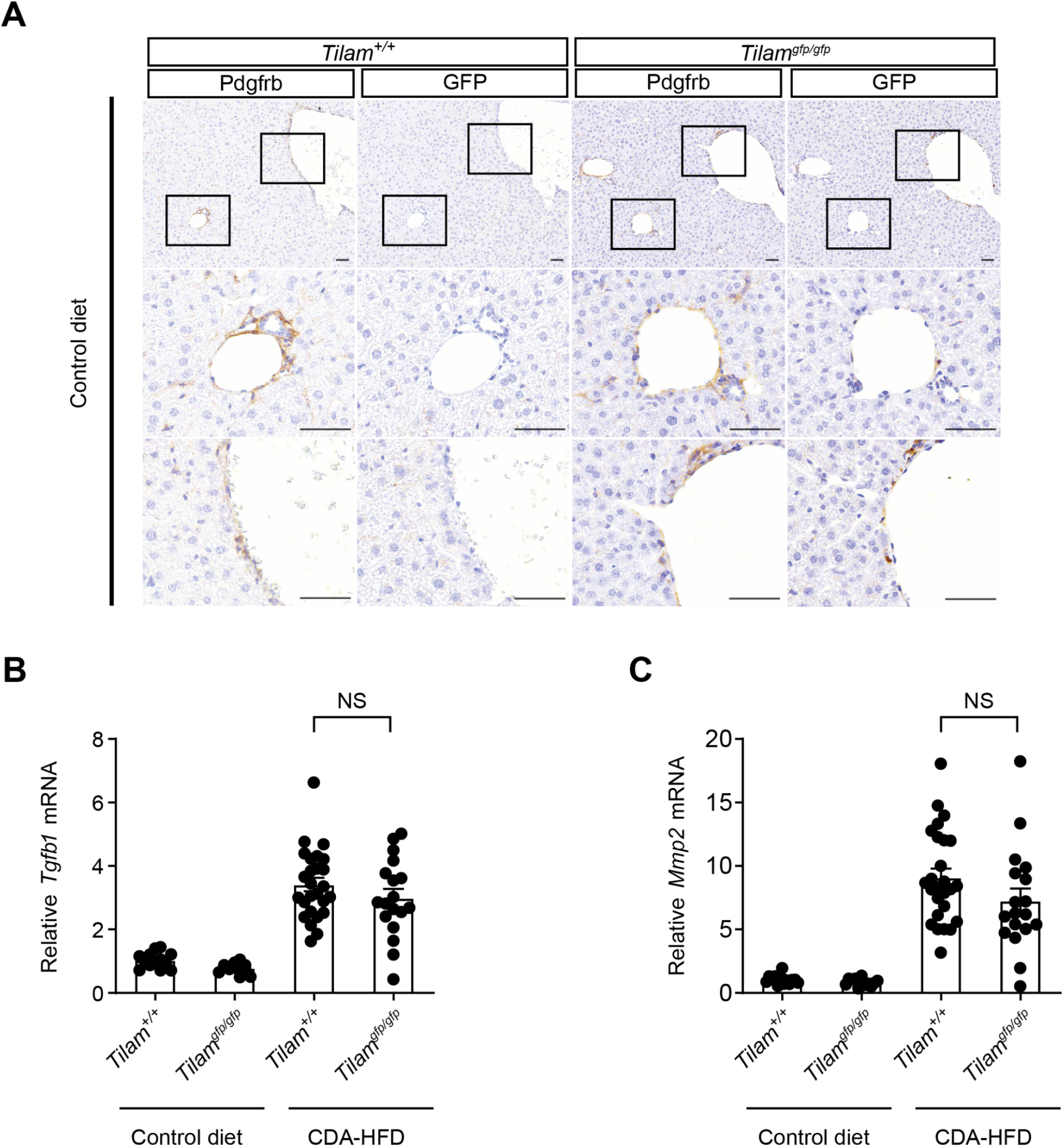
IHC for GFP in control conditions, and *Tgfb1* and *Mmp2* expression in *Tilam^gfp/gfp^*and *Tilam^+/+^* female mice receiving CDA-HFD. (*A*) IHC of Pdgfrb and GFP in *Tilam^+/+^* and *Tilam^gfp/gfp^* livers with normal diet. Pdgfrb is limited to the periportal area and vessels in normal diet and a small number of GFP positive cells are detected in similar regions of *Tilam^gfp/gfp^*mice with control diet. Scale bar: 50 μm. (*B*) *Tgfb1* expression was quantified by qRT-PCR for bulk liver (n=13, 12, 26, 18). Error bars represent mean ± SEM. ns indicates p>0.05 (2-tailed unpaired *t* test). (*C*) *Mmp2* expression was quantified by qRT-PCR for bulk liver (n=13, 12, 26, 18). ns indicates p>0.05 (2-tailed unpaired *t* test).

**Supplementary Figure 6.**
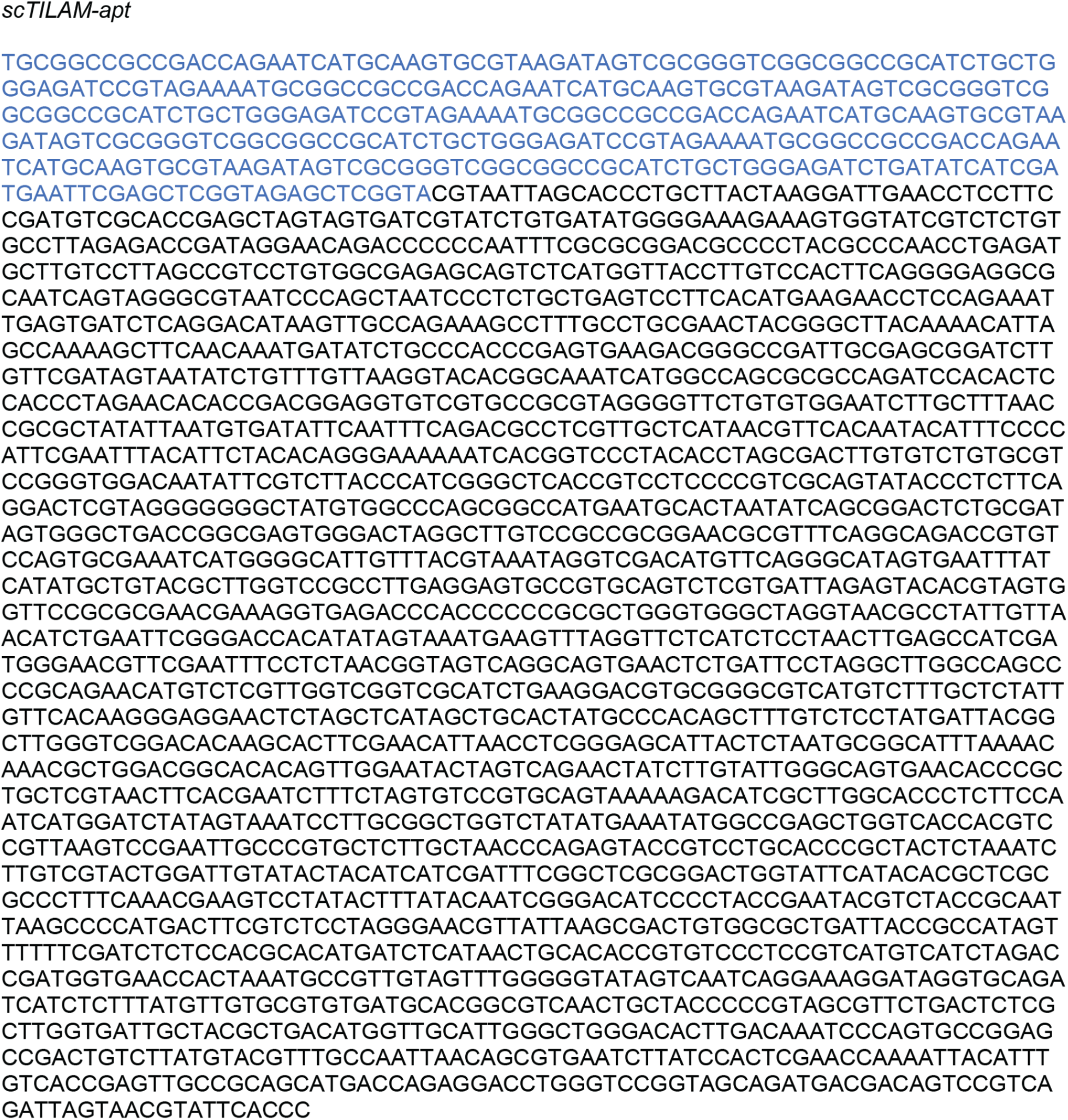
Sequence of scrambled *TILAM* with aptamer (*scTILAM-apt*). The nucleotide sequence of *TILAM was scrambled* and fused to the 4 x S1m aptamer sequence^4, 5^. The aptamer sequence is shown in blue, and the *scTILAM* is shown in black. *TILAM*-apt contains the same sequence as in Supplementary Figure 1B and is fused to the same aptamer sequence on the 5’ end.

**Supplementary Figure 7.**
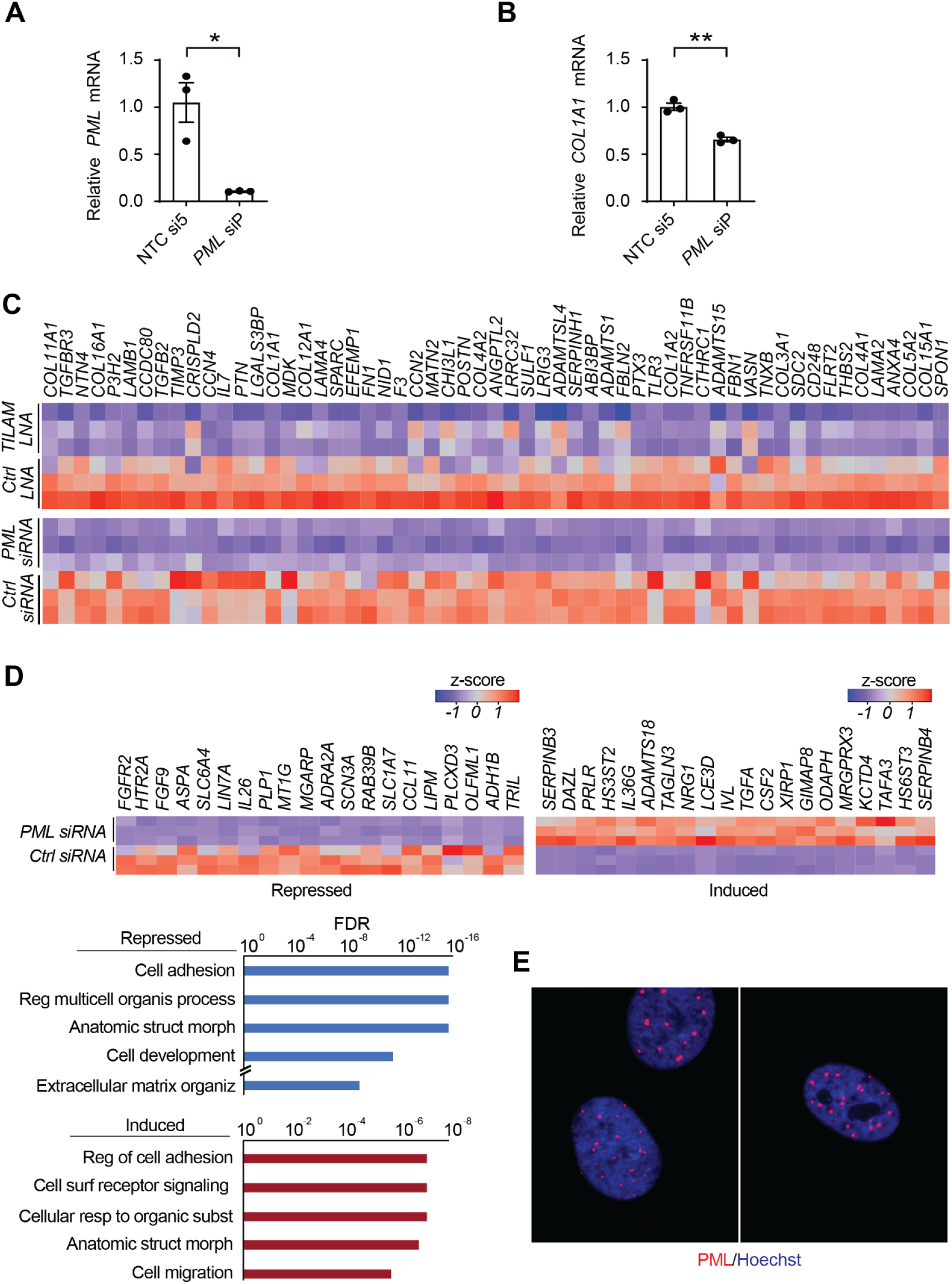
*PML* depletion leads to reduced *COL1A1* expression in primary HSC lines from an additional donor, and depletion leads to broader effects on ECM production as quantified by RNA-seq. (*A*-*B*) *PML* was depleted in primary human HSCs from donor 5 using pooled siRNAs against *PML* mRNA and a control siRNA (NTC 5). *PML* and *COL1A1* levels were quantified by qRT-PCR. * indicates p<0.05, ** indicates p<0.01 (2-tailed unpaired *t* test). Error bars represent mean ± SEM. (*C*) Heatmap shows the relative expression of the 57 genes from the ECM GO category in Figure 7I (GO:0031012) that are repressed by depletion of *TILAM* and *PML*. (*D*) Heatmaps show the relative expression of the top 20 genes repressed (left) and induced (right) with depletion of *PML*. GO analysis was performed on genes repressed (blue) and induced (red) with depletion of *PML*. False discovery rate (FDR) is indicated on the x-axis. (*E*) IF was performed in primary human HSCs using an antibody against PML (red). Nuclei are stained with Hoechst (blue). Three nuclei probed for PML are shown.

## Materials and methods

### Animal studies

All mouse experiments were approved by the IACUC of the Massachusetts General Hospital (2017000074) or Columbia University (AC-AAAF7452). For the carbon tetrachloride (CCl_4_) model of fibrosis, mice received 40% CCl_4_ diluted in olive oil or olive oil control by oral gavage (100 ul total volume) three times a week for four weeks^1^. For the choline-deficient, L-amino acid-defined, high-fat diet (CDA-HFD) model, mice were fed CDA-HFD chow consisting of 60% kcal fat and 0.1% methionine or control chow for twelve weeks^2, 3^. HSCs were sorted from *Lrat-Cre* mice crossed with ZsGreen Cre reporter mice (*R26R^zsGreen^*, Jackson Laboratory) as described^4, 5^ either before or after receiving CCl_4_ by intraperitoneal injections (0.5 ml/g, dissolved in corn oil at a ratio of 1:3) three times a week for four weeks.

### Generation of Tilam mouse

*Tilam^gfp/gfp^* mice were generated by injecting a cocktail of 0.61 pmol/ml each of crRNA+tracrRNA, 100 ng/ml Cas9 protein^6^, and 10ng/ml of donor DNA into pronuclei of E0.5 embryos (C57BL/6). Two crRNAs targeting *Tilam* were mixed 1:1 for injection (Supplementary Table 7). Donor DNA consisted of a plasmid containing GFP and polyA cDNA sequence inserted into exon 1 of *Tilam* (between nt 55 and 56), with 4 kb of genomic homology sequence flanking each side of the insertion site. Post-injection embryos were re-implanted into recipient CD1 pseudo-pregnant females and allowed to develop to term. Pups were screened by genomic PCR to identify founders, initially using a 5’ primer recognizing a sequence outside the 4 kb homology arm and 3’ primer recognizing a sequence in the GFP cDNA (Supplementary Table 7). Founders were back-crossed to C57BL/6 mice (Charles River) to confirm germline transmission and backcrossed for six generations to wild-type C57BL/6 mice before in vivo fibrosis experiments were performed. The locus was amplified and sequenced to confirm that the *GFP-polyA* cassette was inserted between nucleotides 55 and 56 of exon 1 (Supplementary Figure 4C). Subsequent genotyping was performed with primers described in Supplementary Table 7 and through probe sets designed by Transnetyx. Studies were initiated on age-matched wild-type and *Tilam^gfp/gfp^* mice on the same C57BL/6 background at 8-10 weeks of age.

### ARRIVE guidelines

#### Experimental mouse cohort

Both male and female mice were evaluated separately to analyze the phenotype in wild-type and *Tilam^gfp/gfp^* mice. Figure legends indicate the sample size for each result, represented by individual data points. Animal numbers for sample harvest are: wild-type = 135, *Tilam^gfp/gfp^* = 127. Age-matched control wild-type mice of the same C57BL/6 background (Charles River) were purchased and were co-housed for 2 weeks prior to experiments. The mice were fed with the LabDiet Prolab IsoPro RMH 3000, 5P76 and housed in Allentown PIV cage on sani chip hardwood bedding and given carefresh as a nesting material.

#### Cell culture

Primary human HSCs were purchased from Lonza, and LX-2 cells were a gift from Scott Friedman. Cells were cultured as described previously for HSCs^7^. Details for each Individual donor are listed below. Primary mouse HSCs were isolated and cultured as previously described^8^.

| Donor | Vendor ID | Age | Gender | Race | BMI |
| --- | --- | --- | --- | --- | --- |
| 1 | Lonza, HUCNP, ID: 4105 | 45 | M | Caucasian | 24.2 |
| 2 | Lonza, HUCNP, ID: 4270 | 35 | M | Caucasian | 42.1 |
| 3 | Lonza, HUCNP, ID: 4119 | 30 | F | African<br>American | 33.2 |
| 4 | Lonza, HUCLS, ID: 201951 | 28 | M | Caucasian | 24.4 |
| 5 | Lonza, HUCLS, ID: 182821 | 24 | F | African<br>American | 48.8 |

#### Human embryonic stem cell culture (hESCs) and liver organoid differentiation

H1 hESCs (WA01, NIH registration number 0043) were obtained from WiCell, and ESCRO approval for their use was received from UMass Chan Medical School and Massachusetts General Hospital. hESCs were cultured, and genome editing was performed as previously described^9^. In brief, hESCs were co-transfected with Cas9 protein-crRNA-tracrRNA complex (Supplementary Table 7) and homology vector followed by puromycin selection. Individual puromycin-resistant colonies were selected and expanded. Genomic sequencing was then performed to confirm homozygous insertion.

hESCs were differentiated into liver organoids as previously described^10, 11^. In brief, cells were differentiated to definitive endoderm then to foregut spheroids. On day 7 of differentiation, cells were embedded in Matrigel (Corning, 354230) and treated for 4 days with retinoic acid (Tocris, 0695) before switching to Hepatocyte Culture Medium (Lonza, CC-3198). On day 21, liver organoids were extracted from Matrigel and cultured on an orbital shaker^11^. Isolated liver organoids were either treated with TGF-β (10 ng/mL) and FGF-1 (at indicated dose) for 4 days to induce fibrosis or treated with LNAs (500 nM) for 48 hours to deplete *TILAM* expression.

#### Delivery of short interfering (si) RNAs and locked nucleic acids (LNAs)

Human HSCs were reverse transfected with siRNAs^7^, and human HSCs were transduced with LNAs with reverse transfection or nucleofection, as indicated. Reverse transfection was performed using Dharmafect-1 transfection reagent (Horizon Discovery, cat# T-2001) according to the manufacturer’s instructions. For 12-well plates, 60µL of 3µM siRNAs or LNAs were added to 180µL Opti-MEM (Gibco, cat# 31985070) for the final concentration of 150nM and then mixed with diluted Dharmafect-1 in Opti-MEM (2.4 µL Dharmafect-1 in 237.6µL Opti-MEM). After 30 min, HSCs resuspended in transfection medium (DMEM supplemented with 16% FBS) were seeded in wells containing the siRNA or LNA/Dharmafect-1 mixture at ∼70,000 cells/mL in 720µL/well. Transfection in other plate formats were scaled up or down based on surface area. Cells were incubated with siRNAs or LNAs and transfection reagents for 72 hours before analysis. The siRNAs were purchased from Horizon Discovery. and the LNAs were purchased from Qiagen (Supplementary Table 7).

Nucleofection was performed for human HSCs (Figure 1D) and murine HSCs (Figure 4B). 0.5×10^6^of HSCs were resuspended in 1M nucleofection buffer (5 mM KCl, 15 mM MgCl2, 120 mM Na2HPO4/NaH2PO4 pH 7.2, 50 mM Mannitol) and transfected 200 nM LNA/ASO in a Nucleofector 4D (Lonza). HSCs were harvested after 48 hours of treatment. The ASOs against murine *Tilam* were purchased from Integrated DNA Technologies (Supplementary Table 7)

#### qRT-PCR analysis

RNA samples were extracted using Qiagen RNeasy Mini Kit (74104). Using 500 ng total RNA as input, reverse transcription was performed with the iScript gDNA Clear cDNA Synthesis Kit (BIO-RAD, 1725035) according to manufacturer’s instructions. TaqMan Universal PCR Master Mix (Applied Biosystems, cat# 4305719) and TaqMan Real-time PCR Assays (ThermoFisher Scientific) were used for the quantitative real-time PCR (qRT-PCR) analysis of the cDNA samples. *GAPDH* was used as an endogenous control for all experiments unless otherwise stated. The gene-specific Real-time PCR Assays and primers used in this study are listed in Supplementary Table 7.

#### Single molecule RNA fluorescent in situ hybridization (smFISH) and immunofluorescent staining

Single molecule RNA FISH was performed as previously described^12^. Custom Stellaris® FISH probes (Biosearch Technologies) were designed against *TILAM* and *PDGRFB* using the Stellaris® FISH Probe Designer (www.biosearchtech.com/stellarisdesigner, Supplementary Table 8). HSCs were cultured on glass coverslips before fixation and overnight hybridization with Stellaris FISH probe sets labeled with Quasar570 or Quasar670, following the manufacturer’s instructions. (www.biosearchtech.com/stellarisprotocols). Nuclei were defined by Hoechst staining. Imaging was performed with a Nikon 80i upright fluorescence microscope with Hamamatsu Orca CCD camera.

Immunofluorescent staining was performed as previously described^9^ following the sequential immunostaining and single-molecule RNA-FISH protocol for HSCs cultured on glass coverslips. Cells were fixed in 3.7% formaldehyde in DPBS and permeabilized in 1% Triton X-100. Cells were stained with Hoechst and mounted onto slides. Slides were imaged on a StellarVision inverted microscope (Optical Biosystems, model SV20HT) using a Nikon CFI S Plan Fluor ELWD 0.45NA 20XC air objective.

#### Formaldehyde-assisted isolation of regulatory elements (FAIRE)

FAIRE was performed as previously described^12^, and primers are described in Supplementary Table 7.

#### Rapid amplification of cDNA ends (RACE)

Total RNA was isolated from human and murine HSCs. RNA was isolated by the microFast Track 2.0 kit (Life Technologies). The GeneRacer 2.0 kit (Life Technologies) was used to determine the 5’ and 3’ ends of the lncRNA transcripts. Nested 5’ and 3’ RT-PCR was performed in human HSCs and a single round of 5’ and 3’ RT-PCR was performed in murine HCS. Primers are listed in Supplementary Table 7.

#### Nuclear and cytoplasmic fractionation

Nuclear and cytoplasmic fractionation was performed as previously described^12^. Primers used for qRT-PCR are listed in Supplementary Table 7.

#### RNA-Seq

Human HSCs from donor 4 (Lonza, HUCLS, ID: 201951) were transfected with NTC si5 (control), *PML* si17, LNA Ctrl (control), and *TILAM* LNA1 in triplicate. Cells were harvested, and RNA was extracted using Qiagen RNeasy Mini Kit (74104). Analyzed samples showed an RNA quality number (RQN) greater than 9 (Agilent 4200 TapeStation System). PolyA-selection and stranded library preparation (NEB Directional Ultra II RNA library preparation kit) was performed prior to 150 nt paired-end sequencing on a HiSeq4000. Murine HSC libraries were created from sorted HSCs and prepared with the Illumina Truseq mRNA Stranded Prep Kit prior to 100 nt paired-end sequencing on a HiSeq2500.

#### RNA-seq analysis and differential gene expression

For analysis of human HSC data, reads were quality assessed using the FASTQC (v 0.11.9) and aligned to the human reference genome (GRCh38_release_37) from GENCODE with Star aligner (v2.7.10a) using RSEM (v1.3.3) with default parameters. First, the human reference genome was indexed using the GENCODE annotations (gencode.vGRCh38_release_43) with rsem-prepare-reference from RSEM software. Next, rsem-calculate-expression was used to align the reads and quantify gene abundance. The output of rsem-calculate-expression gives separately the read count and transcripts per million (TPM) value for each gene. Differential expression analysis was performed using gene read counts with DESeq2 package (v 1.38.3) to produce LFC values and corresponding p-values (FDR) applying a Benjamini–Hochberg correction for multiple testing, with a minimum of 5 reads required for a differentially-expressed gene. The heatmap was created using normalized gene count values from Deseq2, using R gplots package heatmap.2 function with row scaling. GO analysis was performed and visualized with WebGestalt^13^. Gene expression across cell types and tissues for human and mouse samples were performed as previously described^12, 14^, except that for Supplementary Figure 1C, the original FPKM values were converted into transformed Z-scores so that the values for *Col1a1* and *TIlam* could be compared on the same plot.

#### Identification of orthologous long noncoding RNAs between humans and mice

RNA-seq data from murine HSCs were mapped to the mouse reference genome (mm10), and *ab initio* assembly was performed as previously described^14^. To identify orthologous lncRNAs between humans and mice, we integrated three distinct methods: synteny analysis, whole genome alignment (WGA), and sequence similarity (SS).

1. Synteny: An lncRNA was considered as synteny-orthologous between human and mouse HSCs if it met the following criteria: a) it was expressed in both human and mouse HSCs, b) it was located within 10 kb of the same coding gene (as defined by genes in the human and mouse genomes with the same name), and c) it exhibited consistent relative strand information corresponding to the coding gene.
2. Whole Genome Alignment (WGA): Orthologous lncRNAs identified via WGA were located in the same region of the aligned genomes of humans and mice. We obtained whole-genome alignment chains between the human genome (hg19) and mouse genome (mm10) from the UCSC Genome Browser. We then assigned a location for each lncRNA into 2,000 bins within these aligned chains in both species.
3. Sequence Similarity (SS): An lncRNA was deemed orthologous via sequence similarity if its human and mouse transcripts showed significant similarity, defined by an e-value of less than 1e-5 in a bi-directional search. The sequence alignment search was performed using the BLAST algorithm with the following parameters: blastall -p blastn -i query_seq.fa -d target_seq.fa -e 0.00001 -m 0 -W 8.

#### Hepatic hydroxyproline

To quantify collagen level, liver samples were isolated from the same region of the left liver lobe as described^15^. Isolated samples were homogenized and processed to evaluate hydroxyproline concentration using hydroxyproline assay kits following manufacturer’s instructions (Sigma-Aldrich, MAK008).

#### Collagen proportionate area (CPA)

CPA was measured as described previously^16^. Liver samples were fixed in 4% paraformaldehyde (PFA). Pico-Sirius red staining was performed using the left liver lobe from PFA fixed paraffin embedded sections. Whole sections were scanned and loaded into ImageJ to calculate the ratio of collagen positive area against the total parenchyma area and expressed as a percentage.

#### Immunohistochemistry

Immunohistochemistry (IHC) was conducted on paraffin sectioned samples. For paraffin-embedded sectioning, livers were fixed in 4% paraformaldehyde at 4°C. After dehydration through graded ethanol and paraffin embedment, samples were sectioned at 5-μm. Standard immunofluorescence procedure with antigen recovery was carried out. The following antibodies were used:

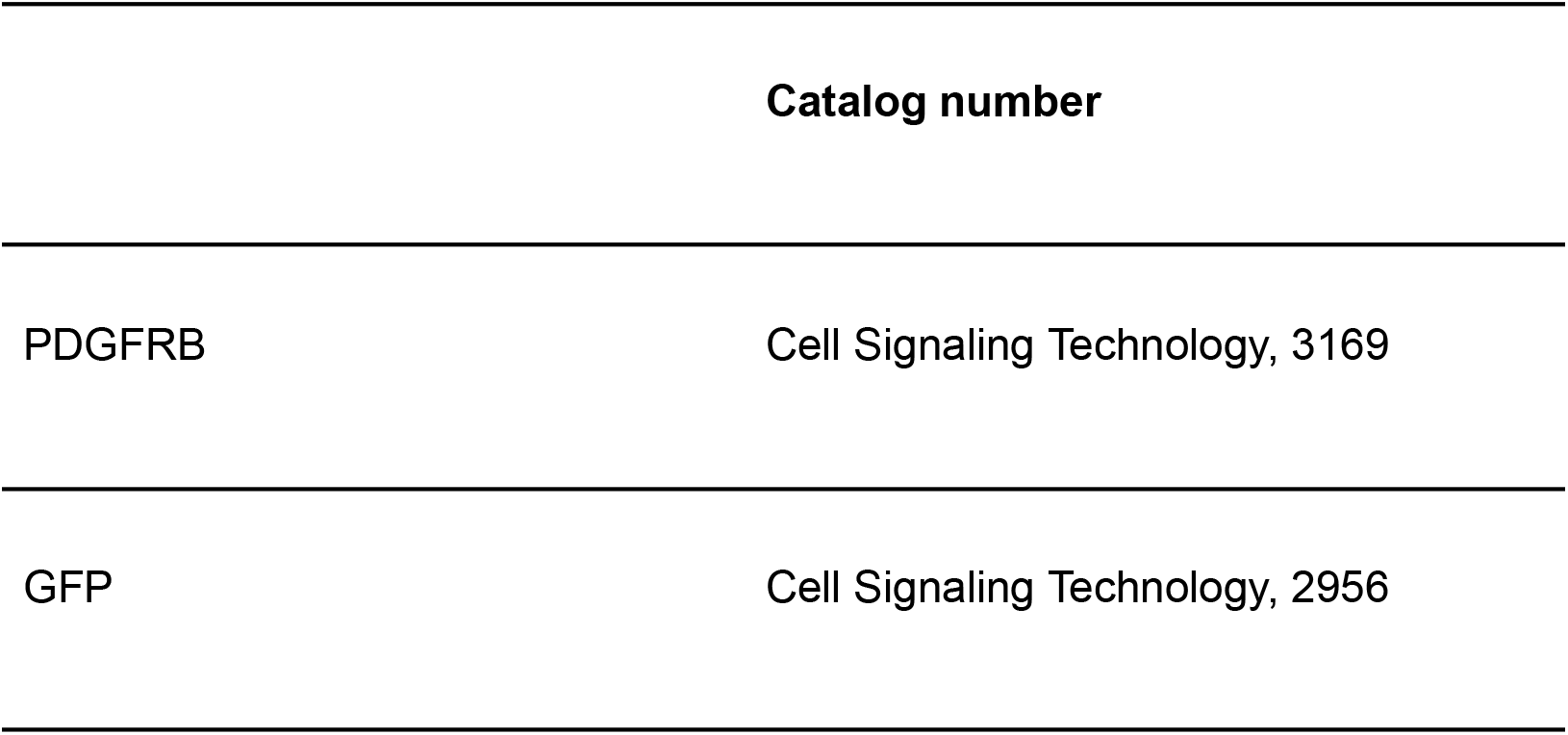

#### Western blotting

Pelleted cells were lysed with RIPA buffer (Thermo Scientific, 89900) supplemented with protease inhibitors (Thermo Scientific, 87786). Cell lysates were centrifuged to remove debris. Protein concentrations were measured using Pierce BCA Protein Assay Kit (Thermo Scientific, 23227). LDS Sample Buffer (Invitrogen, B0007) and Sample Reducing Agent (Invitrogen, B0009) were added to cell lysate, and the sample mixture was boiled for 10 min before loading. 4% to 12% Bis-Tris gels (Invitrogen, NW04120BOX) were used for electrophoresis followed by transfer using iBlot 2 Dry Blotting System (Invitrogen, IB21002S). Membranes were blocked with 1% BSA (Thermo Scientific, 37520) at room temperature for 1 hr and incubated overnight with primary antibody at 4°C. Membranes were washed three times with Tris Buffered Saline-Tween (TBST) buffer (Boston BioProducts, IBB-181–6), incubated with secondary antibody for another 1 hr, washed three times with TBST buffer, and then incubated with SuperSignal West Pico PLUS chemiluminescent substrates (Thermo Scientific, 34580) for 5 min before scanning with a LI-COR Odyssey M imaging system. GAPDH and PML antibodies were used at 1:1000 dilution.

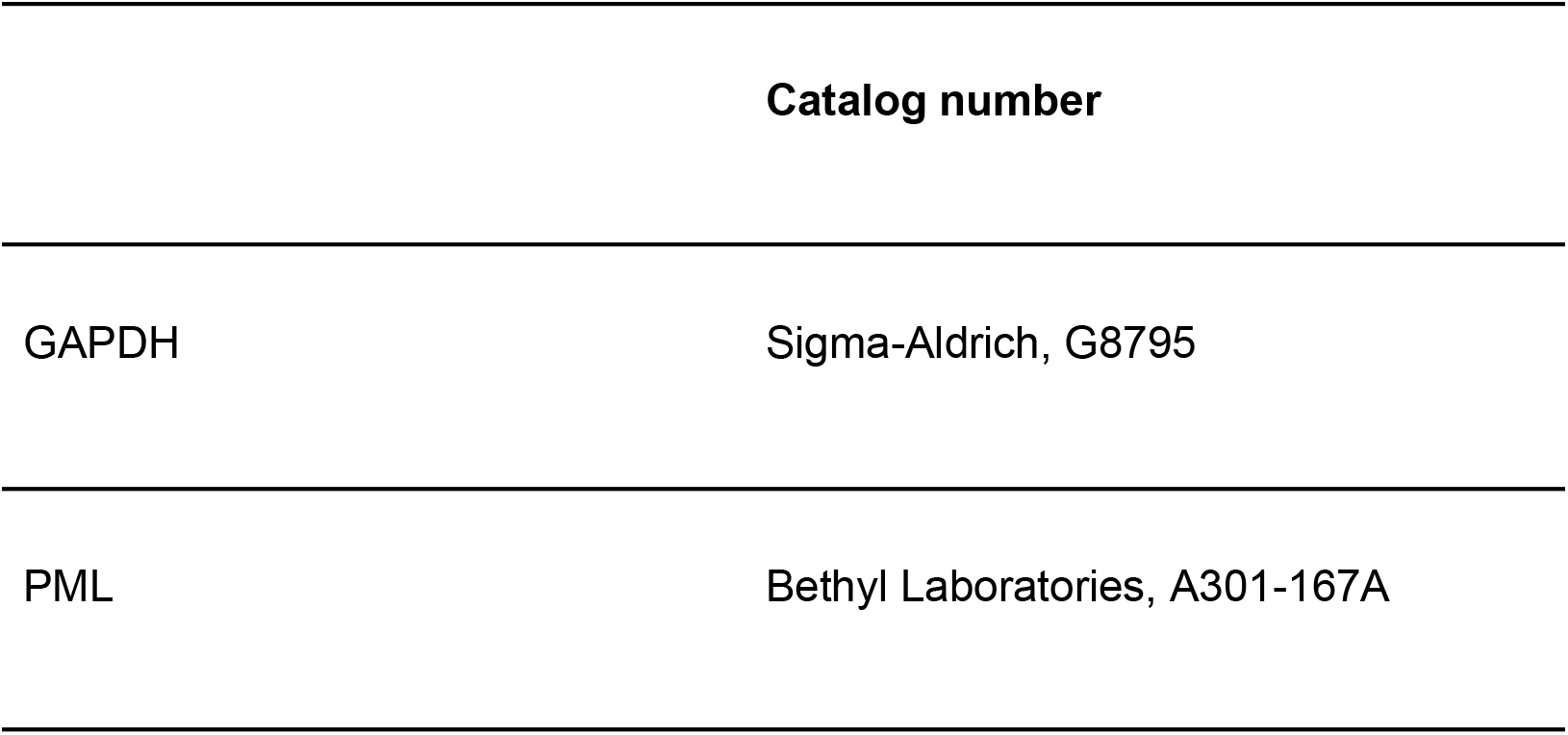

#### Mass spectrometry and analysis

LX-2 cells were infected with lentivirus expressing *TILAM*-*apt* or *SCRM-apt* and selected with puromycin. Confluent cells were rinsed with DPBS and exposed to 400 mJ/cm^2^ of energy on ice using a UV cross-linker (Stratalinker). UV cross-linked cells were lysed (150 mM KCl, 25 mM Tris-HCl pH 7.4, 5 mM EDTA, 5 mM MgCl2, 1% NP-40, 0.5 mM DTT, Roche mini-tablet protease inhibitor, 100 U/mL RNAseOUT), and debris was removed by centrifugation at 16000 g. The resulting lysate was further treated with 50 μL of Avidin Agarose beads (Thermo Fisher, 20219) to eliminate biotin from the lysates. The cleared lysate was then incubated with 150 μL of Streptavidin C Dynabeads (Thermo Fisher) on a rocker at 4°C for 4 hours. The beads were collected using a magnet and washed three times with a wash buffer similar to the lysis buffer, except for an increased KCl concentration of 350 mM. To release the proteins, the beads were incubated with RNase A and resuspended in 2x LDS sample buffer. Lysates were resolved on a polyacrylamide gel, and segments ranging from 30 to 350 kDa were excised for subsequent mass spectrometry analysis. The resulting proteomic data were analyzed using Crapome proteomic analysis (https://reprint-apms.org/) with a threshold of >90% confidence and a greater than four-fold enrichment over controls.

